# Pretrainable Geometric Graph Neural Network for Antibody Affinity Maturation

**DOI:** 10.1101/2023.08.10.552845

**Authors:** Huiyu Cai, Zuobai Zhang, Mingkai Wang, Bozitao Zhong, Quanxiao Li, Yuxuan Zhong, Yanling Wu, Tianlei Ying, Jian Tang

**Affiliations:** BioGeometry, Office 1505, 8 Haidianbeierjie St., Beijing, 100083, China; Mila - Québec AI Institute, 6666 St-Urbain, Montréal, H2S 3H1, Québec, Canada; Department of Computer Science and Operations Research, Université de Montréal, 2920 chemin de la Tour, Montréal, H3T 1J4, Québec, Canada; Department of Decision Sciences, HEC Montréal, 3000 chemin de la Côte-Sainte-Catherine, Montréal, H3T 2A7, Québec, Canada; Shanghai Engineering Research Center for Synthetic Immunology, Fudan University, 131 Dong’an Street, Shanghai, 200032, China; MOE/NHC/CAMS Key Laboratory of Medical Molecular Virology, Shanghai Frontiers Science Center of Pathogenic Microorganisms and Infection, Shanghai Institute of Infectious Disease and Biosecurity, School of Basic Medical Sciences, Fudan University, 131 Dong’an Street, Shanghai, 200032, China

## Abstract

Increasing the binding affinity of an antibody to its target antigen is a crucial task in antibody therapeutics development. This paper presents a pretrainable geometric graph neural network, GearBind, and explores its potential in *in silico* affinity maturation. Leveraging multi-relational graph construction, multi-level geometric message passing and contrastive pretraining on mass-scale, unlabeled protein structural data, GearBind outperforms previous state-of-the-art approaches on SKEMPI and an independent test set. A powerful ensemble model based on GearBind is then derived and used to successfully enhance the binding of two antibodies with distinct formats and target antigens. ELISA EC_50_ values of the designed antibody mutants are decreased by up to 17 fold, and ***K*_D_** values by up to 6.1 fold. These promising results underscore the utility of geometric deep learning and effective pretraining in macromolecule interaction modeling tasks.

## Introduction

Antibody plays a crucial role in the human immune system and serves as a powerful diagnostic and therapeutic tool, due to its ability to bind selectively and specifically to target antigens with high affinity. *In vivo*, antibodies go through affinity maturation, where the target-binding affinity gradually increases as a result of somatic hypermutation and clonal selection [1]. When a new antigen surfaces, therapeutic antibody leads repurposed from known antibodies or screened from a natural or *de novo* designed library often require *in vitro* affinity maturation to enhance their binding affinity to a desired, usually sub-nanomolar, level.

Wet lab experimental methods for *in vitro* antibody affinity maturation usually involve constructing mutant libraries and screening with display technology [2–5]. These methods, while significantly improved during the past few years, are still labor-intensive and costly in general, taking 2-3 months or more to complete the process. Let’s consider the combinatorial search space of possible mutations. There are usually 50-60 residues on the complementarity-determining region (CDR) of an antibody, which are hypervariable *in vivo* and contribute to the majority of the binding free energy Δ*G*_bind_ [6]. Previous works show that multiple point mutations are often needed for successful affinity maturation [7, 8]. Performing experiments on all combinations of over a thousand possible point mutations in antibody CDR regions (60 residues *×* 19 residues per residue) is difficult if not prohibitive. Therefore, a fast and accurate computational method for narrowing down the search space is much desired.

Nevertheless, it is nontrivial for computational affinity maturation methods to balance speed and accuracy. Molecular dynamics methods based on empirical force fields [9–12] rely on human knowledge and abstractions to evaluate binding free energy changes after mutations. However, accurate models are often too slow to be used for ranking thousands of mutations (let alone their combinations). In recent years, machine learning, and particularly deep learning, has been demonstrated as a powerful tool capable of tackling this dilemma. Many machine learning methods [13–18] formulate the affinity maturation problem as a structure-based binding free energy change (ΔΔ*G*_bind_ := Δ*G*_bind_^(mt)^ *−* Δ*G*_bind_^(wt)^, where *wt* is short for *wild type* and *mt* denotes *mutant*) prediction problem. However, despite the importance of protein side-chain conformation to protein-protein interaction, most existing methods model atom-level geometric information indirectly or incompletely, e.g. using hand-crafted features or residue-level features. These approaches inadequately address the intricate interplay between side-chain atoms. Another critical problem is the massive amount of paired binding affinity data required by machine learning models for them to become accurate and reliable. To the best of our knowledge, the largest publicly available protein-protein binding free energy change dataset, Structural Kinetic and Energetic database of Mutant Protein Interactions (SKEMPI) v2.0 [19], contains only 7085 ΔΔ*G*_bind_ measurements on 348 protein complexes, a tiny amount compared to the training set sizes of foundational protein models such as AlphaFold2 [20] and ESM2 [21].

To tackle the aforementioned challenges, we introduce GearBind, a pretrainable deep neural network that leverages multi-level geometric message passing to model the nuanced protein-protein interactions. We utilize contrastive pretraining techniques on large-scale protein structural dataset to incorporate vital structural insights into the model (Fig. 1). *In silico* experiments on SKEMPI and an independent test set demonstrate the superior performance of GearBind and the benefit of pretraining. We combine the GearBind models with previous state-of-the-art methods to create an ensemble model that achieves state-of-the-art performance on all metrics. Ablation study confirms the importance of key design choices within GearBind and the key role it played in the ensemble. We then use the GearBind-based ensemble to perform *in silico* affinity maturation for two antibodies with distinct formats and target antigens. Binding of the antibody CR3022 against the spike (S) protein of the Omicron SARS-CoV-2 variant is increased by up to 17 fold as measured by Enzyme-linked immunosorbent assay (ELISA), and by 6.1 fold as measured by Bio-layer Interferometry (BLI), after synthesizing and testing only 20 antibody candidates. All designed antibodies have maintained or increased binding towards the receptor-binding domains (RBDs) of both the SARS-CoV-2 Delta variant and SARS-CoV. Binding of the fully human single-domain antibody (UdAb) against the oncofetal antigen 5T4 is increased by up to 5.6 fold as measured by ELISA, and by up to 2.1 fold as measured by BLI, after testing 12 candidates. These results underscore the importance of geometric deep learning and effective pretraining on antibody affinity maturation and, more generally, macromolecule interaction modeling.

**Fig. 1:**
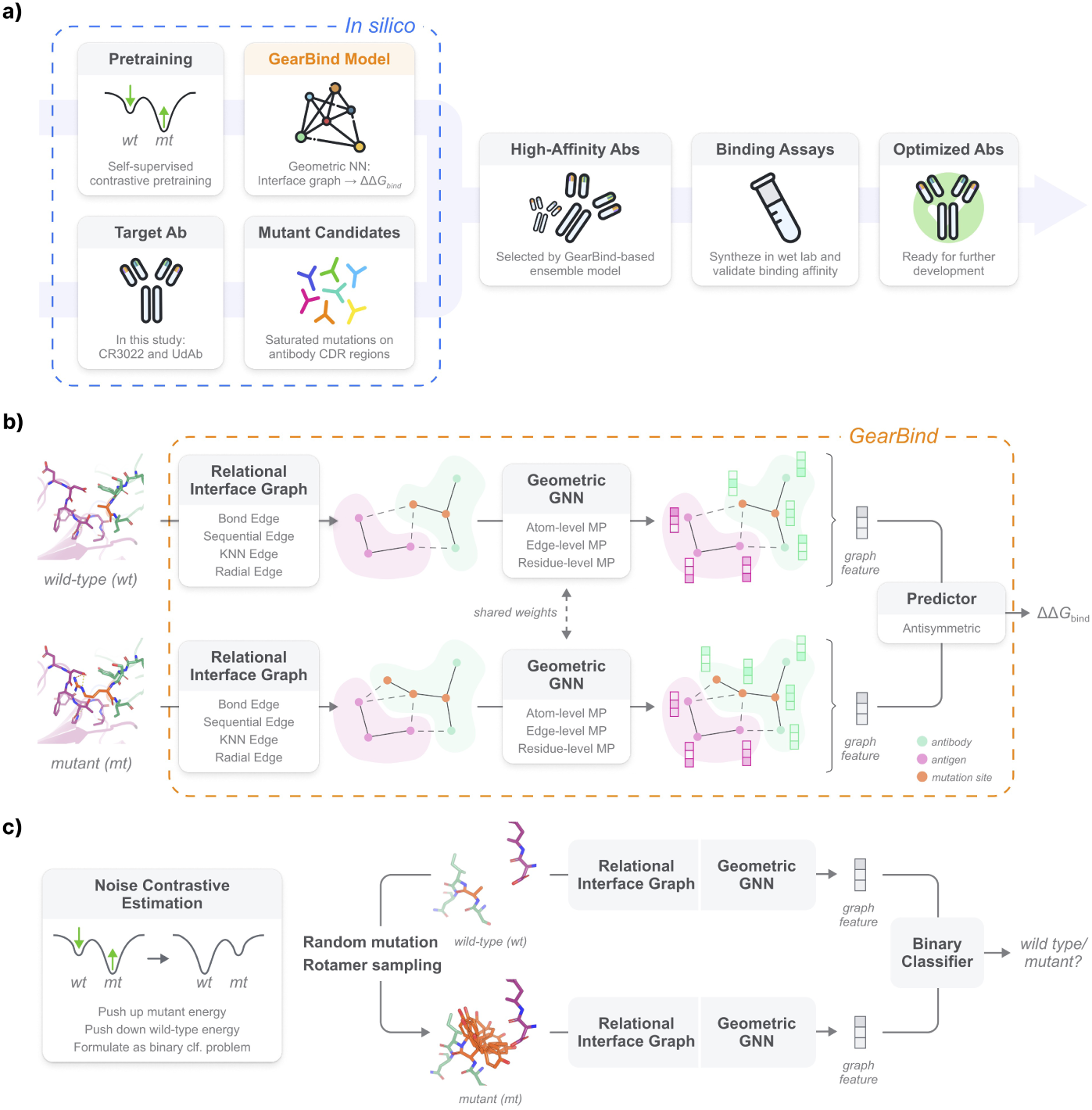
GearBind-based *in silico* antibody affinity maturation pipeline. (a) Pipeline Overview: The pipeline features the geometric neural encoder, GearBind, which undergoes self-supervised pretraining on CATH and supervised learning on SKEMPIv2. The GearBind-based ensemble model is employed to perform *in silico* affinity maturation on a target antibody, given its bound structure to the native antigen. Guided by the model predictions, antibodies with improved binding affinity can be found after testing one or two dozen mutant candidates. NN: neural network. Ab: Antibody. CDR: Complementarity-determining region. Designed using resources from Flaticon.com. (b) GearBind Model: GearBind employs a shared graph neural network to encode both the wild-type and mutant complex structures. For each structure, a relational interface graph is constructed. A geometric graph neural network, GearNet, then performs multi-relational and multilevel message passing on the graph to extract rich interface representations. The mutational effect ΔΔ*G*_bind_ is predicted by an antisymmetric predictor given the GearNet-extracted representations of the two complexes. (c) Selfsupervised Pretraining: GearBind+P leverages mass-scale unlabeled protein structures via self-supervised, contrastive pretraining. The model is trained to contrast between native structures and randomly mutated structures with side-chain torsion angles sampled from a rotamer library. Pretraining helps GearBind+P explore the energy landscape of native protein structures and results in improved performance in ΔΔ*G*_bind_ prediction.

## Results

### GearBind: a pretrainable ΔΔ*G*_bind_ predictor

The GearBind framework is designed to extract geometric representations from wild-type and mutant structures via multi-level and multi-relational message passing to predict the binding free energy change ΔΔ*G*_bind_. GearBind leverages information within a protein complex at three different levels with complementary insights, namely, atom-level information holding precise spatial and chemical characteristics, edge-level information capturing angular relationships, and residue-level information highlighting broader context within the protein complex. Merging these distinct yet interconnected tiers of information allows for a more holistic view of protein complexes, potentially enhancing model capabilities.

More formally, when a protein complex structure is input to GearBind, a multi-relational interface atom graph is first constructed to model the detailed interactions within the complex. The relations defined cover both sequential proximity (for atoms on the same chain) and spatial proximity (which includes *k*-nearest-neighbor and within-*r*-radius relations). Atom-level representations are obtained by applying a geometric relational graph neural network (GearNet [22]) on the interface graph. On top of that, a line graph is constructed by treating each edge in the atom graph as a line node, connecting adjacent line nodes, and encoding the angular information as line edge features. Edge-level interactions are then captured by performing message passing on the line graph, similar to a sparse version of AlphaFold’s triangle attention [20]. Finally, after aggregating atom and edge representations for each residue, a geometric graph attention layer is applied to pass messages between residues. This *multi-level message passing* scheme injects multi-granularity structural information into the learned representations, making it highly useful for the task of ΔΔ*G*_bind_ prediction.

Although GearBind can be trained from scratch on labeled ΔΔ*G*_bind_ datasets, it could suffer from overfitting or poor generalization if the training data size is limited. To address this problem, we propose a self-supervised pre-training task to exploit large-scale unlabeled protein structures in CATH [22, 23]. In the pretraining stage, the encoder is trained to model the distribution of the native protein structures via noise contrastive estimation [24]. Specifically, we maximize the probability (*i.e.* push down the energy) of native CATH proteins while minimizing the probability of mutant structures (Fig. 1c) generated by randomly mutating residues and sampling side-chain torsional angles from a rotamer library [25]. Distinguishing native, stable protein structures from sampled mutant structures pushes the model towards understanding side-chain interaction patterns, which are crucial to protein-protein binding. Through this process, meaningful knowledge from abundant single-chain protein structural data could be transferred to benefit protein-protein binding modelling.

### Cross Validation on SKEMPI

We validated GearBind performance via a split-by-complex, five-fold cross validation on SKEMPI v2.0. Our splitting strategy dictates that each test set share no common PDB complex with its corresponding training set, making it more realistic than the split-by-mutation strategy, where the wild-type protein complexes and even the mutation sites in the test set could appear during training. We compared GearBind and GearBind+P (pretrained GearBind fine-tuned on SKEMPI) to state-of-the-art physics-based tools FoldX [9], Flex-ddG [10] and the deep learning method Bind-ddG [8]. The results (Table 1) show that GearBind, with its multi-relational graph construction and multi-level message passing schemes, outperforms the baselines in terms of mean absolute error (MAE), root mean squared error (RMSE) and PearsonR, while seconds FoldX in terms of SpearmanR. Pretraining GearBind brings further performance improvements, resulting in +5.4% SpearmanR, +2.6% PearsonR, *−*2.4% MAE and *−*1.7% RMSE. This highlights the effective knowledge transfer from mass-scale, unlabelled protein structural data.

**Table 1:** Cross validation performance of different methods on SKEMPI (*n* = 5729). For each metric, we report the mean and standard error of the mean. “+P” means “with geometric pretraining on CATH”. MAE: Mean average error. RMSE: root mean square error. Among individual models, the best and the second-best performing model for each metric is highlighted in bold and italic, respectively.

| Model | MAE↓ | RMSE↓ | PearsonR↑ | SpearmanR↑ |
| --- | --- | --- | --- | --- |
| FoldX [9] | 1.364 ± 0.134 | 2.027 ± 0.170 | 0.491 ± 0.007 | **0.526 ± 0.011** |
| Flex-ddG [10] | 1.236 ± 0.101 | 1.849 ± 0.150 | 0.497 ± 0.034 | 0.484 ± 0.020 |
| Bind-ddG [8] | 1.255 ± 0.096 | 1.759 ± 0.125 | 0.581 ± 0.037 | 0.443 ± 0.041 |
| GearBind | <u>1.143 ± 0.088</u> | <u>1.639 ± 0.103</u> | 0.659 ± 0.030 | 0.498 ± 0.033 |
| GearBind+P | **1.115 ± 0.072** | **1.611 ± 0.075** | **0.676 ± 0.041** | <u>0.525 ± 0.046</u> |
| Ensemble | **1.028 ± 0.080** | **1.503 ± 0.101** | **0.729 ± 0.016** | **0.643 ± 0.030** |

To understand the contributions of key architectural design choices in GearBind, we benchmarked the performance of 5 GearBind variants on SKEMPI. As shown in 2e, the tested GearBind variants perform worse than GearBind on all four metrics. The exclusion of edge- and residue-level message passing from GearBind brings a 13% and 3% SpearmanR drop, highlighting the benefits of combining multi-level information during feature extraction. The exclusion of side-chain atoms from the interface graph hurts performance even more (15% SpearmanR drop), showing the importance of explicitly modeling the full-atom structure. Notably, replacing the multi-relational interface graph with a KNN graph results in a severe 23% SpearmanR decline, while training a simple RGCN model on the multi-relational graphs results in performance on par with Bind-ddG (*−*9% SpearmanR compared to GearBind, +2% compared to Bind-ddG). This suggests that the multi-relational graph construction strategy is a key ingredient in GearBind.

### A GearBind-based ensemble for in silico affinity maturation

To understand the behavior of the benchmarked models on SKEMPI, we binned the SKEMPI dataset by the target difficulty and plotted the PearsonR and SpearmanR of each model on targets with different difficulty levels. PDB codes in SKEMPI are categorized into “easy” (50+ similar data points in training set), “medium” (1-50), and “hard” (0) targets based on the number of training data points having a high structural similarity (TM-score [26] *>* 0.8) to it. Deep learning models, namely Bind-ddG, GearBind and GearBind+P, enjoys superior performance compared to physics-based methods such as FoldX and Flex-ddG on easy targets, but the table turns when we move to the hard targets (Fig. 2a,b), showing room for improvement in their generalization capabilities. We also studied the performance on mutations that cause low (*<* 0.5 kcal/mol), medium (0.5 *−* 2) and high (*>* 2) absolute changes in binding free energy. As Fig. S9 shows, all models perform better when the binding level changes more drastically. GearBind achieves outstanding performance in this region, with a PearsonR value of 0.707, compared to FoldX’s 0.411, showing its potential to identify mutations that could significantly enhance or disrupt binding. When the *|*ΔΔ*G*_bind_*|* is small, predictions from all methods have very low correlation with experimental ΔΔ*G*_bind_ values, hinting either the noises in data or a deficiency of current tools in modeling weaker, more intricate interactions.

**Fig. 2:**
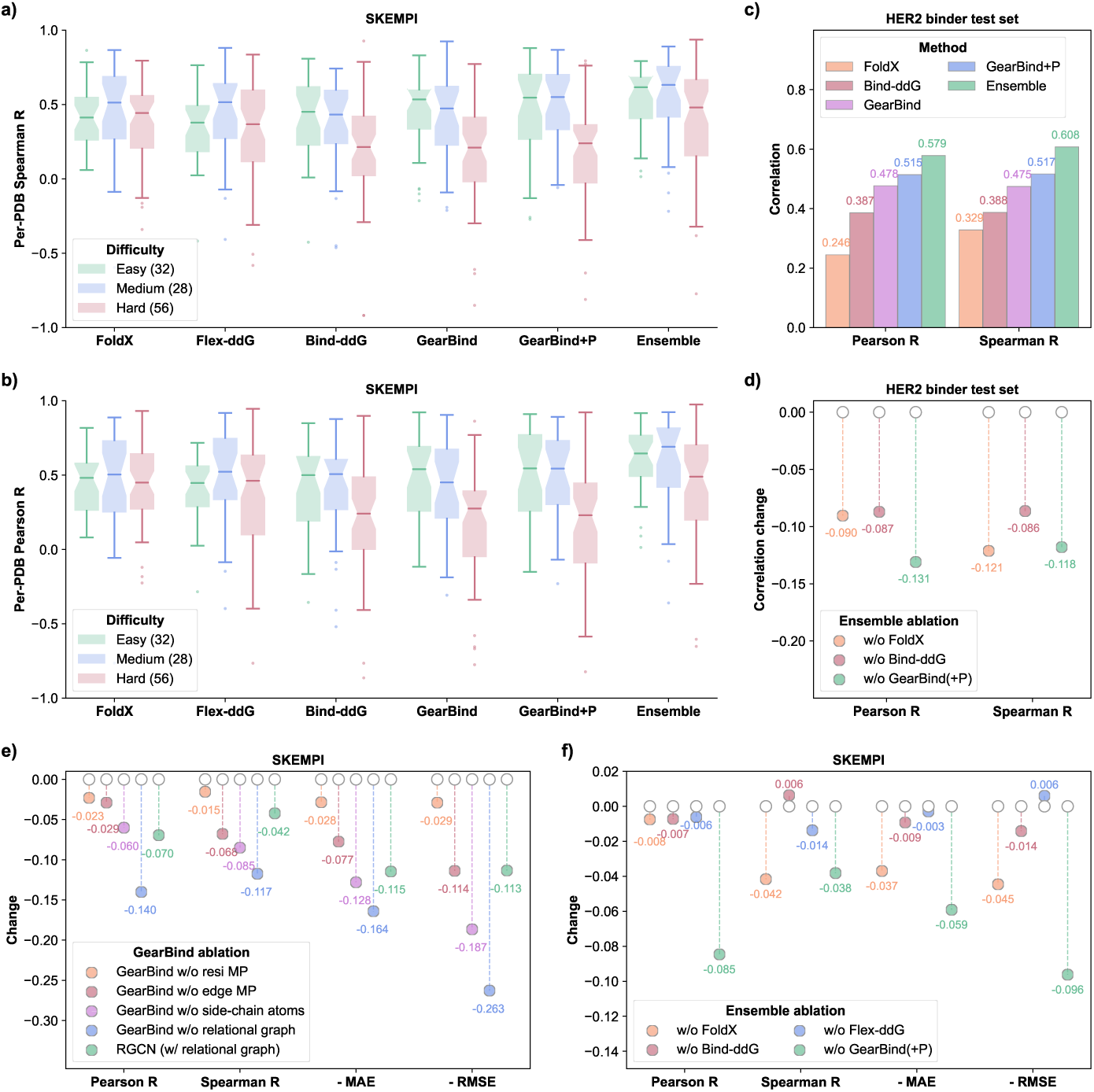
*In silico* evaluation on SKEMPI and the HER2 binders test set. Comparative analysis of Per-PDB Spearman (a) and Pearson (b) correlations between predictions of various models and experimental data across SKEMPI subsets with varying difficulty levels. PDB codes in SKEMPI are categorized into “easy” (50+ similar data points in training set), “medium” (1-50), and “hard” (0) targets based on the number of training data points having a high structural similarity (TM-score *>* 0.8) to it. The number of PDB codes for each difficulty is annotated in the figure legends. The box spans the interquartile range (25th to 75th percentile), with a solid line inside marking the median. Outliers are determined by 1*×* inter-quartile range. (c) Benchmark results on the HER2 binders test set (*n* = 419) show Pearson and Spearman correlations for various models. The deep learning models are trained on SKEMPI. (d) Change of performance metrics in HER2 binders test set when excluding various models from the FoldX + Bind-ddG + GearBind(+P) ensemble. (e) Change of performance when changing GearBind architecture design, as quantified by cross validation performance on SKEMPI (*n* = 5729). (f) Change of performance on SKEMPI (*n* = 5729) when excluding different models from the FoldX + Flex-ddG + Bind-ddG + GearBind(+P) ensemble.

To combine the advantages of both physics-based and deep learning methods, we used the ensemble of all benchmarked methods to perform subsequent *in silico* affinity maturation. The prediction of the ensemble model is the simple average of prediction values from FoldX, Flex-ddG, GearBind, GearBind+P and Bind-ddG. The proposed ensemble model outperforms each individual model in all four evaluation metrics (Table 1). We evaluated the contribution of individual models to the ensemble by excluding each of them and evaluating performance on SKEMPI. The results (Fig. 2f) show that excluding GearBind and GearBind+P hurts overall performance the most. Specifically, for the PearsonR metric, the exclusion of FoldX, Flex-ddG and Bind-ddG individually results in a marginal (less than 0.01) decrease, but the removal of GearBind causes a significant (more than 0.08) decline. We also note that, while FoldX is not the best-performing model when used in isolation, removing it from the ensemble results in the biggest SpearmanR drop. This shows that FoldX plays an important role in complementing the deep learning models and forging a robust and accurate ensemble model. In fact, combining GearBind, GearBind+P and FoldX yields comparable performance to the 5-model ensemble (Fig. S26).

### Evaluation on the HER2 binders test set

With the models built and trained (on SKEMPI), we tested their performance on the HER2 binders test set, which we collected from [27]. This dataset contains high-quality binding affinity data, measured by surface plasmon resonance (SPR) on 419 HER2 binders with *de novo* designed CDR loops. The antibodies in the dataset are variants of Trastuzumab that have high edit distance (7.6 on average), making them potentially challenging for ΔΔ*G*_bind_ predictors trained on low-edit-distance data. Among the benchmarked methods (Flex-ddG is not benchmarked due to its high time cost), GearBind+P achieve the best PearsonR and SpearmanR (Fig. 2c). We then averaged the predictions of all benchmarked models to form an ensemble model, and excluded each model from the ensemble to measure the performance change. Similarly, excluding GearBind(+P) hurts PearsonR the most, and excluding FoldX hurts SpearmanR the most, with GearBind(+P) closely following (Fig. 2d).

### Affinity Maturation of CR3022 and anti-5T4 UdAb

To validate the efficacy of our methodology, two antibodies, CR3022 and anti-5T4 UdAb, were selected as subjects for affinity maturation. The CR3022 antibody, originally isolated from a convalescent SARS patient [28], has been subsequently identified to bind to SARS-CoV-2 [29, 30]. Meanwhile, a UdAb directed against the oncofetal antigen 5T4, is characterized by its exceptional stability [31]. Note that the two antibodies are in distinct formats and target distinct antigens. Both antigens have only one structurally similar protein chain (TM-score *>* 0.8) with a different binding site in SKEMPI, making them challenging targets for our pipeline (Table S9, S10, Fig. S14, S15).

For CR3022, a total of 12 mutants were picked out in the first-round experimental validation according to the ensemble prediction of their binding affinity changes against the RBDs of the wild-type, BA.1.1, and BA.4 SARS-CoV-2 strains. We note that the wild-type and Delta RBDs share the same amino acids at the interface to CR3022. In an ELISA pre-experiment, we tested the binding of these mutants to the RBD of the SARS-CoV-2 Delta strain with antigen concentration at 100 nM. Nine out of twelve candidates exhibited improved binding compared the wild-type CR3022 (Fig. S16a). In the further validation with reduced RBD concentration at 10 nM, the EC_50_ values for all 9 candidates were lower than the wild-type CR3022 (Fig. 3a, b). Based on these results, we combined the well-performed CR3022 mutations and synthesized 8 candidates with double or triple mutations as our second-round designs. Seven of the eight designed multi-point mutants exhibited enhanced binding against the Delta RBD, with 1.8 to 3.4 fold lower ELISA EC_50_ compared to the wild-type. The triple mutant SH100D+SH103Y+SL33R has the lowest EC_50_ at 0.06 nM (Fig. 3c, d). Against the Omicron Spike protein, the above seven multi-point mutants again displayed 7.6 to 17.0 fold binding increase with sub-nanomolar EC_50_ values, among which the SH100D+SH103Y+SL33R triple mutant still exhibited the best performance (Fig. 3e, f). We next tested the binding of the mutants designed in the second-round against the RBD of SARS-CoV, to examine if binding optimization of CR3022 against SARS-CoV-2 RBDs caused a drastic change in binding to its original target. Seven out of eight mutants did not exhibit a significant change in ELISA EC_50_against SARS-CoV RBD (Fig. 3g, h). In summary, the above results demonstrate the success of our GearBind-based pipeline in CR3022 antibody affinity optimization.

**Fig. 3:**
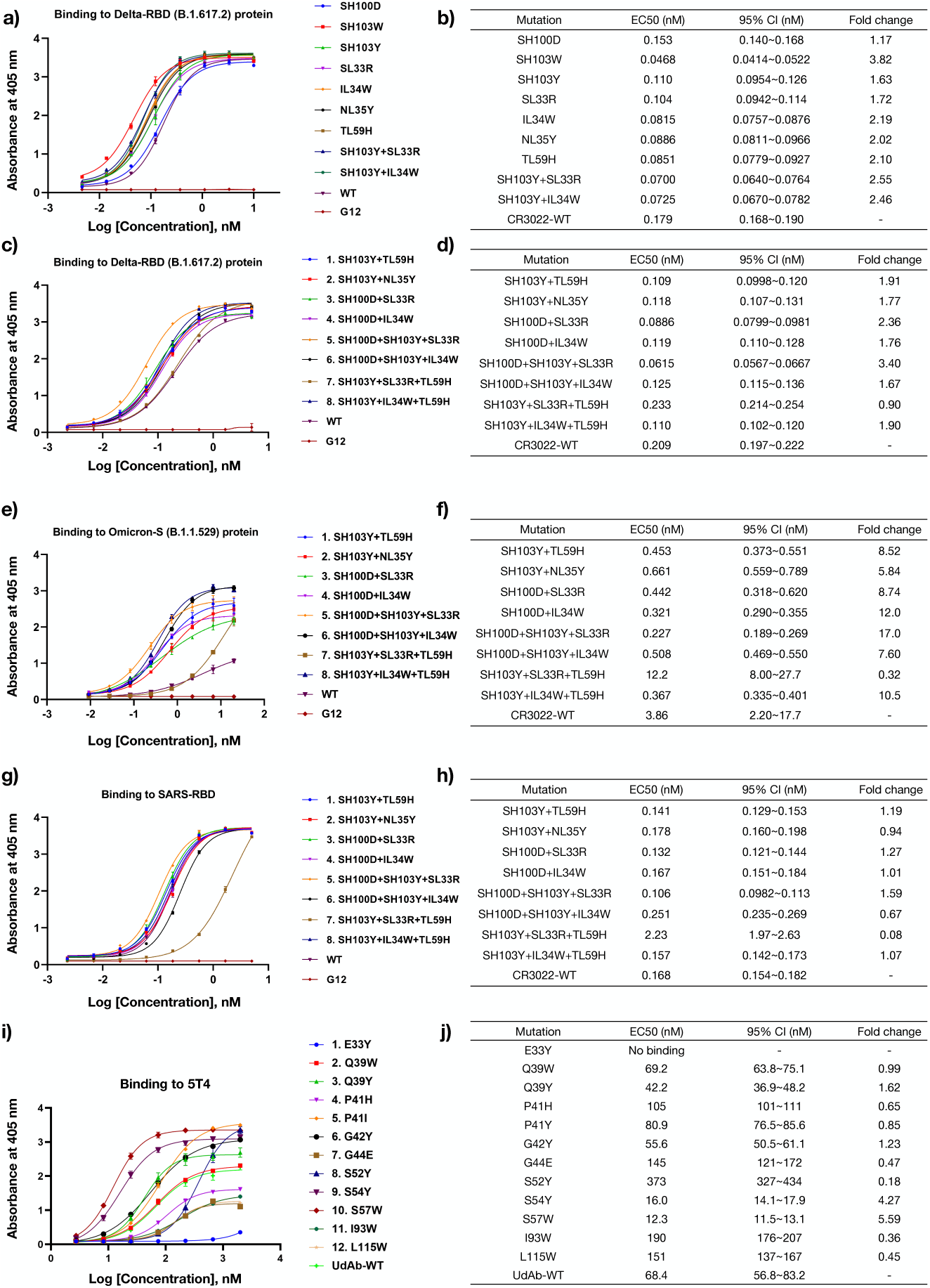
ELISA binding assay results for CR3022 and anti-5T4 UdAb mutants designed with a GearBind-based pipeline. On the left panels (a, c, e, g, i), the concentration-response curves alongside with EC_50_ values evaluated from ELISA assays are displayed, with the center denoting the mean absorbance and error bars indicating the standard deviation from three technical duplicates. On the right panels (b, d, f, h, j), the fitted EC_50_ values, their 95% confidence intervals and the fold changes in binding calculated as 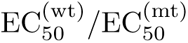 are displayed. Tested systems include the first-round CR3022 designs binding to Delta RBD (a, b); the second-round CR3022 designs binding to Delta RBD (c, d), Omicron S protein (e, f) and SARS-RBD (g, h); and single point mutants of anti-5T4 UdAb (i, j) binding to 5T4.

To demonstrate the generalizability of our method, we extended our experimental validation to the anti-5T4 UdAb. We developed 12 single-point mutants using the GearBind-based pipeline and verified their binding to 5T4 with ELISA. The highest binding UdAb mutant was S57W, with a 5.6 fold decrease in EC_50_. This highlights the potential of our approach in enhancing antibody affinity for antibodies of different formats and with different targets, making it a promising tool for therapeutic antibody development.

We further validated the affinity-matured CR3022 antibodies and anti-5T4 UdAbs using Bio-layer Interferometry (BLI) to assess their binding affinities more accurately (Fig. 4). The 7 tested CR3022 mutants show a 3.1 to 6.1 fold increase in binding affinity against the Omicron Spike protein, with the best performing mutant being SH100D+SH103Y+IL34W. The SH100D+SH103Y+SL33R triple mutant, identified by ELISA as the best-performing mutant, exhibits a 4.1 fold increase in binding affinity. The two tested anti-5T4 UdAb mutants, S54Y and S57W, exhibited a 1.8 fold and 2.1 fold improvement in binding affinity, respectively. Overall, the BLI measurements are consistent with the ELISA binding assay results and demonstrate the increased binding affinity of CR3022 and UdAb variants designed by the GearBind-based pipeline.

**Fig. 4:**
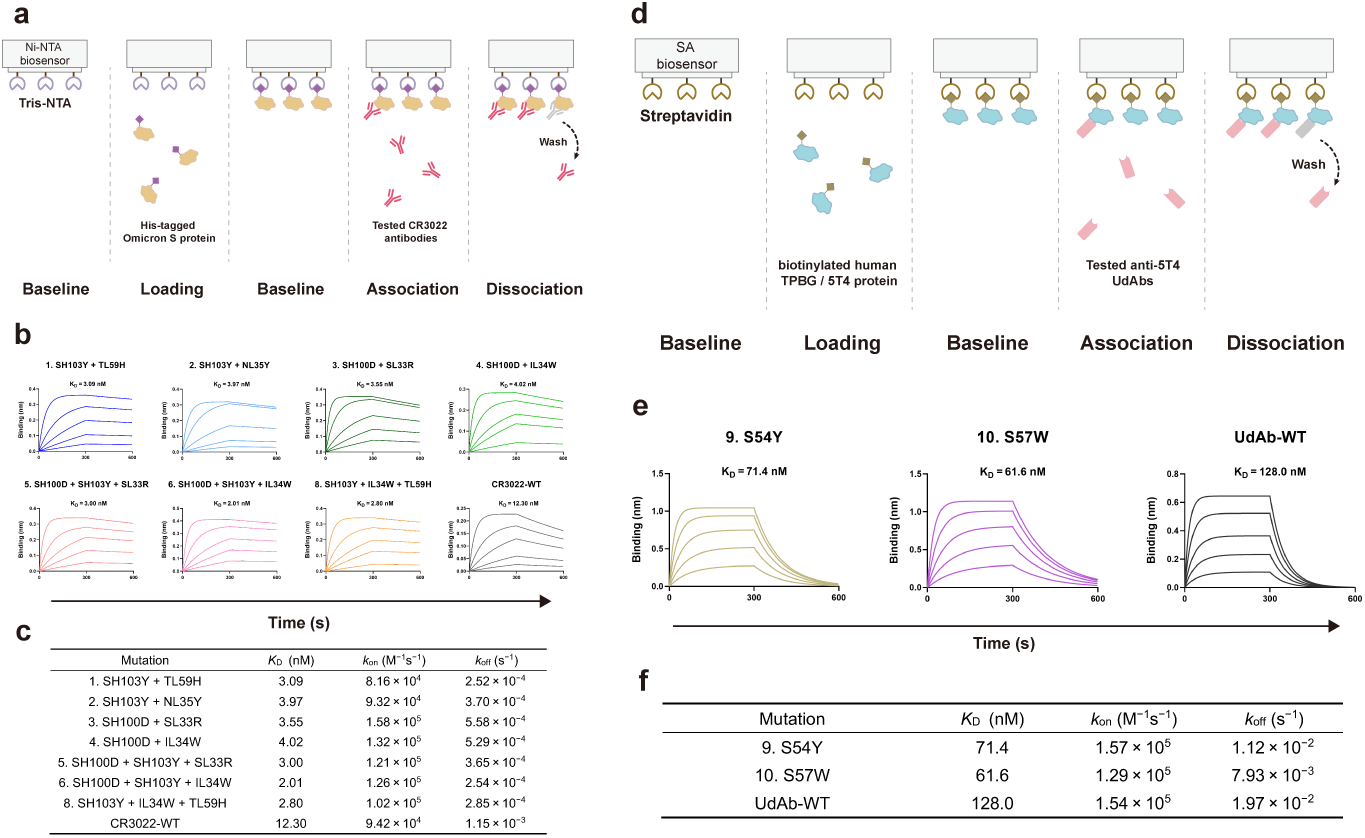
Bio-layer interferometry binding assay for CR3022 and anti-5T4 UdAbs candidates. (a, d) Illustration of the experiment protocol with (a) Ni-NTA and (d) SA biosensors. (b, e) The binding kinetics of different CR3022 (b) and anti-5T4 UdAbs (e) candidates. The starting concentration for each antibody was 300 nM. The data were determined by fitting curves to a global 1:1 binding model. The *K*_D_ (equilibrium dissociation constant) values, annotated on each plot, were determined with *R*^2^ values of greater than 98% confidence level. (c, f) The *k*_on_ (association rate constant), *k*_off_ (dissociation rate constant), and *K*_D_ values of mutant and wild-type CR3022 antibodies (c) and anti-5T4 UdAbs (f).

### Structural Characteristics of Optimized Antibodies

Understanding the sequence-structure-function relationship of mutations designed by deep learning not only helps improve our models but also aids in interpreting their biological significance. To explore the underlying principles governing the increased antibody-antigen binding, we carried out molecular dynamics simulations and structural analyses on both the wild-type and mutant antibodies with the lowest ELISA EC_50_, namely, the SH100D+SH103Y+SL33R triple mutant for CR3022, and the S57W mutant for the anti-5T4 UdAb. We conducted a 1 µs all-atom molecular dynamics simulation at room temperature for each system, including their respective wild-type counterparts (see Methods for details).

Based on the simulation results, among the four mutations studied, three demonstrated an increased number of hydrogen bonds with the target: SH100D and SL33R from CR3022, and S57W from UdAb (Fig. 5g, h). These four mutations in CR3022 and UdAb also stabilized the antibodies, as shown by the reduced fluctuations in C*α* atoms in the corresponding antibody chains (Fig. S20). Although the S103Y mutation in the heavy chain of CR3022 did not increase polar contacts, it potentially enhanced hydrophobic interactions between the antibody and antigen by excluding more solvent due to the larger size of tyrosine. In summary, the mutations designed by our pipeline likely achieved higher binding affinity through the formation of new interactions, while we also observed stabilized binding residues and altered structural properties in the mutated structures.

**Fig. 5:**
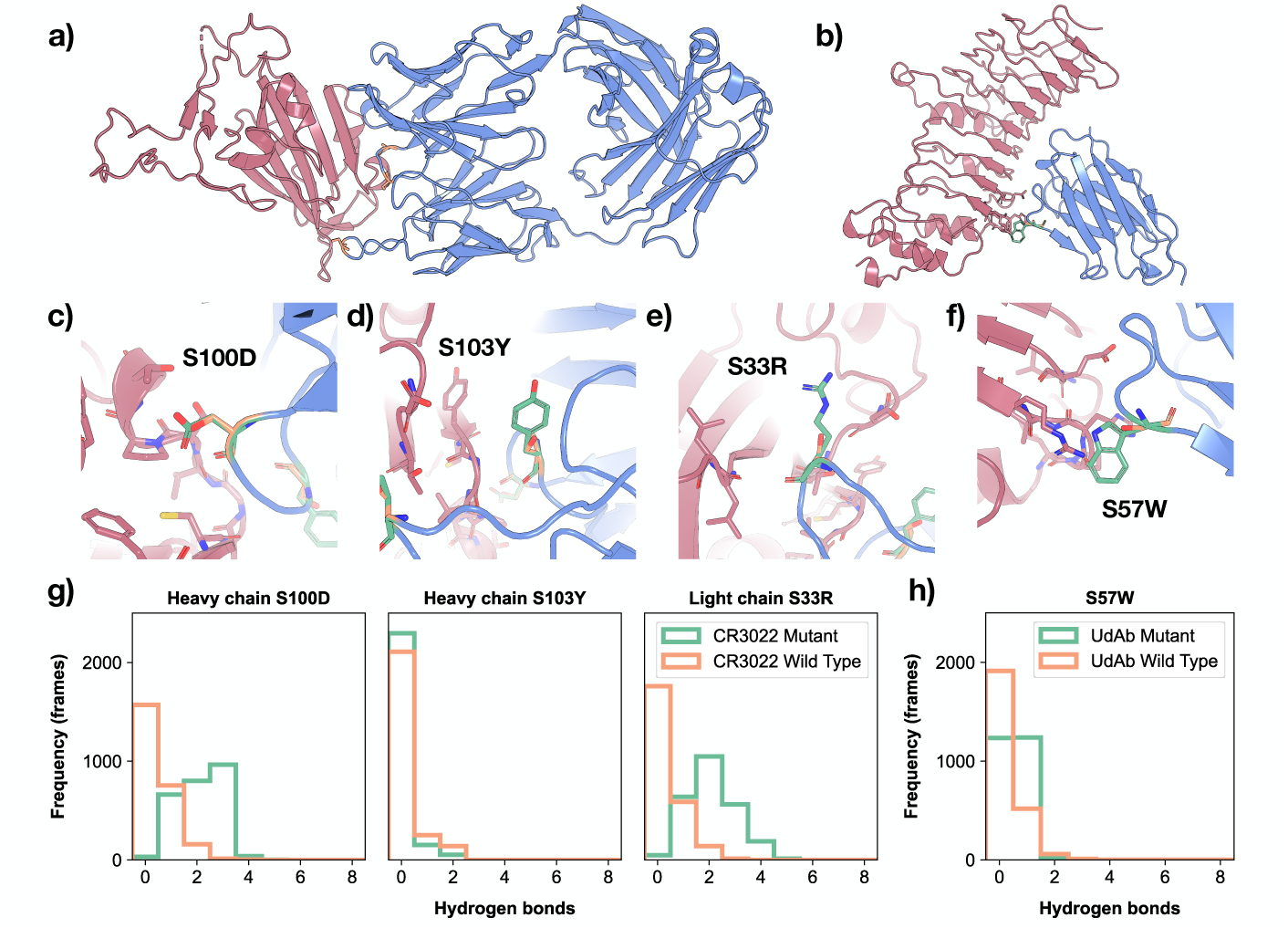
Structural analysis of optimized CR3022 and anti-5T4 UdAbs. (a) Complex structure of antibody CR3022 and the SARS-CoV-2 RBD used for affinity maturation. The target antigen is colored in red, and the antibody in blue. Mutation sites S100D, S103Y in heavy chain and S33R in light chain are marked in orange. (b) Complex structure of the single-domain antibody UdAb and its target oncofetal antigen 5T4. Mutation site S57W is marked in orange. (c-e) Three mutation sites in the CR3022 triple-mutant, namely S100D (c), S103Y (d) in heavy chain and S33R (e) in light chain. (f) The S57W mutation site in the UdAb single-point mutant. (g) Depicts the number of hydrogen bonds between the mutation sites and target antigen in the molecular dynamics simulation of CR3022 and RBD complex. (h) Depicts the number of hydrogen bonds between mutation sites and target antigen in the molecular dynamics simulation of UdAb and 5T4 complex. In (g,h), hydrogen bond distributions from wild types are colored in orange, and from mutants are colored in green.

Interestingly, GearBind-predicted contribution (Fig. S24) provided further insight into the formation of potential contacts for these designed mutations. Most contributions were found to be consistent with our deductions based on the molecular dynamics simulations. The possible hydrophobic interaction in S103Y was also presented in the contributions, providing further validation to our findings and aligning well with our deductions based on protein structure (Fig. S24c).

## Discussion

This study reports an *in silico* antibody maturation pipeline based on a pretrainable geometric graph neural network, GearBind, and the successful application of the pipeline on two distinct antibodies, CR3022 and anti-5T4 UdAb. Substantial *in silico* experiments were done to evaluate model performance and understand their strengths and limitations. The technical strengths of the proposed GearBind model can be summarized as follows: (1) In the graph construction phase, a multi-relational graph is built upon all heavy atoms on the interface. The relations defined cover both sequential proximity and spatial proximity. Replacing the all-atom graph to backbone-atom-only graph, or replacing the multi-relational graph to a simple kNN graph both cause severe performance decline. (2) In the feature extraction phase, a multi-level message passing scheme is employed to obtain a comprehensive view on the intricate interactions at protein interfaces. (3) A unique pretraining algorithm based on contrastive learning is proposed, which harnesses the abundant, unlabeled single-chain protein structures in CATH, distills knowledge about side-chain torsion angles into the model to further boost its performance.

We challenged our GearBind-based pipeline with two real-world antibody affinity maturation projects. ELISA binding assays showed that CR3022 mutations proposed by the pipeline have led to successful enhancements in binding against both the Delta RBD and the Omicron Spike protein. Notably, 7 out of 10 CR3022 single-point mutants and 9 out of 10 multi-point mutants showed a significant increase in binding, with up to 3.8 fold decrease in ELISA EC_50_ for the Delta RBD and 17.0 fold for the Omicron Spike protein. Among 12 single-point anti-5T4 UdAb mutants, our pipeline achieved a maximum decrease of 5.6 times in ELISA EC_50_ and 2.1 times in BLI-measured *K*_D_. In short, GearBind proves to be an efficient and powerful tool for the design of antibodies with enhanced binding affinities. Based on the molecular dynamics simulations of the top-performing mutants identified by the GearBind-based pipeline, we observed that our designs enhanced binding affinity by creating new interactions or strengthening existing contacts, particularly hydrogen bonds. This provides insight into how GearBind learns from data and designs mutations that increase binding affinity.

While we mainly focus on structure-based methods in this work, others have explored purely sequence-based models for affinity maturation [32]. Our evaluation of ESM-1b and ESM-1v models on SKEMPI (Table S4 and Fig. S12) results in negative SpearmanR values, hinting that zero-shot prediction of large-scale protein language models is not a generally reliable method for ranking the binding affinity of protein complexes [33]. This result is reasonable because the “fitness” of a peptide sequence, as modelled by protein language models, does not necessarily imply strong binding to all other biomolecules. For example, improved fitness of the Spike protein of SARS-CoV-2 would likely involve decreased binding affinity towards existing neutralizing antibodies. Another argument is that structural information plays a key role in building an accurate and reliable algorithm for protein-protein interactions [34].

Looking ahead, the potential applications of GearBind reach beyond protein-protein binding optimization. The model can be readily adapted to tackle protein-peptide and protein-ligand docking challenges, thereby opening up possibilities for its use in minibinder and enzyme design.

Despite these positive prospects, we acknowledge certain limitations in our current methodology and discuss potential directions for future work. Firstly, the prerequisite for structure-based ΔΔ*G*_bind_ prediction is an accurate complex structure, which is not easily available for most antibody-antigen pairs. To address this problem, homology modeling tools [35] can be used to build the complex structure from a template structure. This is how we built the complex structure of CR3022 binding to the Omicron RBDs. A more aggressive approach is to directly predict the complex structure from the sequence. As multimer structure prediction methods become more and more accurate [36], they might one day become reliable as the starting point of structure-based affinity maturation. Secondly, the reliance on external tools for mutant structure generation increases the time cost, and limits our action space to substitutions only. Future efforts can focus on training end-to-end models that directly predict the ΔΔ*G*_bind_, and models that can account for amino acid insertion and deletion. We also call for better pretraining strategy and architecture design to improve the generalization capabilities of deep learning models, making them more robust on proteins they have not seen before. All in all, we believe our work takes a solid step towards building a reliable, robust and efficient *in silico* affinity maturation pipeline that would bring tremendous opportunities to research and drug discovery applications.

## Methods

### Datasets

#### SKEMPI

We used the SKEMPI v2 [19] dataset for training and validation. The dataset contains 7,085 ΔΔ*G*_bind_ measurements on 348 complexes. We performed pre-processing following [13, 18], discarding data with ambiguous ΔΔ*G*_bind_ values (e.g. the mutant is a non-binder without exact *K*_D_ measurements) or high ΔΔ*G*_bind_ variance across multiple measurements (*>* 1 kcal/mol), and arrived at 5,729 distinct mutations, with their ΔΔ*G*_bind_ measurements, on 340 complexes. See Table S6 for a list of discarded, high-variance SKEMPI mutations. For each mutation, we sampled the mutant structure with FoldX 4 [9] based on the wild-type crystal structure. We used PDBFixer v1.8 [37] to fix the PDB structures beforehand if the raw structure could not be processed by FoldX. These FoldX-derived SKEMPI structures are used to train deep learning models, including Bind-ddG, GearBind and GearBind+P. The resulting dataset was split into five subsets with roughly the same size using a split-by-PDB strategy, in order to perform cross validation.

#### CATH

For pretraining, we use a non-redundant subset of CATH v4.3.0 domains, which contains 30,948 experimental protein structures with less than 40% sequence identity. We also remove proteins that exceed 2,000 AAs in length for efficiency. During pretraining, we randomly truncate long sequences into 150 AAs for efficiency. It is important to note that, currently, our pretraining exclusively utilizes single-chain proteins. The information learned by single-chain pretraining can be transferred to downstream tasks on protein complexes and we have found that this approach alone is sufficient to yield improvement.

#### HER2 binders

The HER2 binders test set was collected from [27]. The raw data include SPR data for 758 binders and 1097 non-binders. As all benchmarked methods only support amino acid substitutions, we filter out the binders that have different lengths compared to the wild-type antibody (Trastuzumab), leaving 419 Trastuzumab mutants. ΔΔ*G*_bind_ values are calculated by 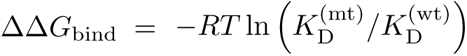 based on the SPR-measured binding affinity. Note that we only use this dataset as a test set to evaluate physics-based models (FoldX, Flex-ddG) and deep learning models (Bind-ddG, GearBind and GearBind+P) trained on SKEMPI.

### GearBind implementation

Given a pair of wild-type and mutant structures, GearBind predicts the binding free energy change ΔΔ*G*_bind_ by building a geometric encoder on a multi-relational graph, which is further enhanced by self-supervised pre-training. Note that the key feature that makes the neural network *geometric* is that it considers the spatial relationship between entities, *i.e.*, nodes in a graph. In the following sections, we will discuss the construction of multi-relational graphs, multi-level message passing and pre-training methods.

### Constructing relational graphs for protein complex structures

Given a protein-protein complex, we construct a multi-relational graph for its interface and discard all other atoms. Here a residue is considered on the interface if its Euclidean distance to the nearest residue from the binding partner is no more than 6Å. Each atom on the interface is regarded as a node in the graph. We add three types of edges to represent different interactions between these atoms. For two atoms with a sequential distance lower than 3, we add a **sequential edge** between them, the type of which is determined by their relative position in the protein sequence. For two atoms with spatial distance lower than 5Å, we add a **radial edge** between them. Besides, each atom is also linked to its **10-nearest neighbors** to guarantee the connectivity of the graph. Spatial edges that connect two atoms adjacent in the protein sequence are not interesting and thus discarded. The relational graph is denoted as 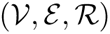 with 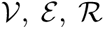 denoting the sets of nodes, edges and relation types, respectively. We use the tuple (*i, j, r*) to denote the edge between atom *i* and *j* with type *r*. We use one-hot vectors of residues types and atom types as node features for each atom and further include sequential and spatial distances in edge features for each edge.

### Building geometric encoder by multi-level message passing

On top of the constructed interface graph, we now perform multi-level message passing to model interactions between connected atoms, edges and residues. We use 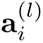 and 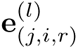 to denote the representations of node *i* and edge (*j, i, r*) at the *l*-th layer. Specially, we use 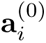 to denote the node feature for atom 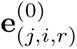 and to denote the edge feature for edge (*j, i, r*). Then, the representations are updated by the following procedures:

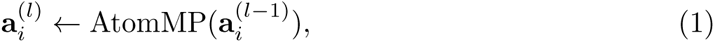

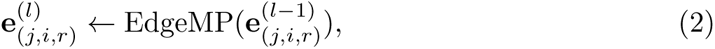

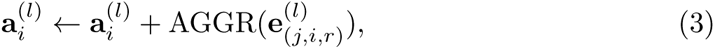

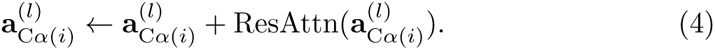

First, we perform atom-level message passing (AtomMP) on the atom graph. Then, a line graph is constructed for the message passing between edges (EdgeMP) so as to learn effective representations between atom pairs. The edge representations are used to update atom representations via an aggregation function (AGGR). Finally, we take the representations 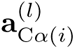 of the alpha carbon as residue representation and perform a residue-level attention mechanism (ResAttn), which can be seen as a special kind of message passing on a fully-connected graph. In the following paragraphs, we will discuss these components in details.

#### Atom-level message passing

Following GearNet [22], we use a relational graph neural network (RGCN) [38] to pass messages between atoms. In a message passing step, each node aggregates messages from its neighbors to update its own representation. The message is computed as the output of a relation (edge type)-specific linear layer when applied to the neighbor representation. Formally, the message passing step is defined as:

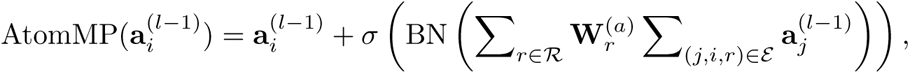

where BN(*·*) denotes batch norm and *σ*(*·*) is the ReLU activation function.

#### Edge-level message passing and aggregation

Modeling sequential proximity or spatial distance alone is not enough for capturing the complex protein-protein interactions (PPI) contributing to binding. Multiple works have demonstrated the benefits of incorporating angular information using edge-level message passing [20, 22, 39]. Here we construct a line graph [40], *i.e.* a relational graph among all edges of the above atom-level graph. Two edges are connected if and only if they share a common end node. The relations, or edge types, are defined as the angle between the atom-level edge pair, discretized into 8 bins. We use 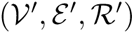 to denote the constructed line graph. Then, relational message passing is used on the line graph:

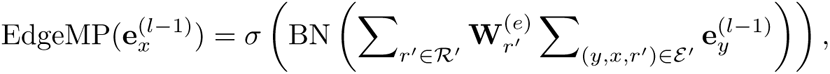

where *x* and *y* denote edge tuples in the original graph for abbreviation.

Once we updated the edge representations, we aggregate them into its end nodes. These representations are fed into a linear layer and multiplied with the edge type-specific kernel matrix 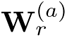 in AtomMP:

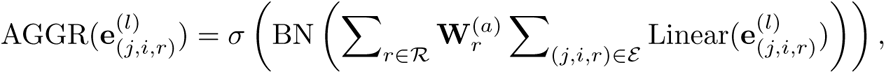

which will be used to update the representation for atom *i* as in equation 3.

#### Residue-level message passing

Constrained by the computational complexity, atom and edge-level message passing only consider sparse interactions while ignoring global interactions between all pairs of residues. By modeling a coarse-grained view of the interface at the residue level, we are able to perform message passing between all pairs of residues. To do this, we design a geometric graph attention mechanism, which takes the representations of the alpha carbon of residues as input and updates their representations with the output as in equation 4. Here we follow the typical definition of self-attention to calculate attention logits with query and key vectors and apply the probability on the value vectors to get residue representations ***r****_i_*:

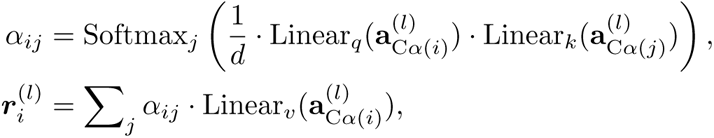

where *d* is the hidden dimension of the representation 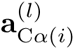 and the Softmax function is taken over all *j*.

Besides traditional self-attention, we also include geometric information in the attention mechanism, which should be invariant to roto-translational transformations on the global complex structure. Therefore, we construct a local frame for each residue with coordinates of its Nitrogen, Carbon and alpha Carbon atoms:

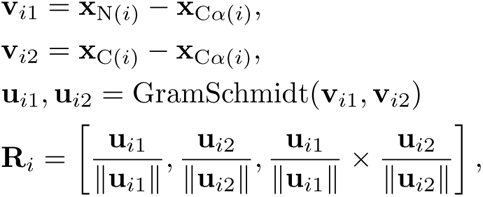

where we use **x** to denote the coordinate of an atom and GramSchmidt(*·*) refers to the Gram–Schmidt process for orthogonalization. Then, the geometric attention is designed to model the relative position of beta carbons of all residues *j* in the local frame of residue *i*:

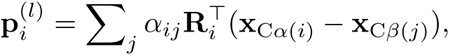

where 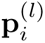 is the spatial representations for the residue *i*. When the complex structure is rotated, the frame **R***_i_* and relative position **x**_C*α*(*i*)_ *−* **x**_C*β*(*j*)_ are rotated accordingly and the effects will be counteracted, which guarantees the rotation invariance of our model.

The final output is the concatenation of residue representations 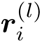 and spatial representations 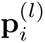:

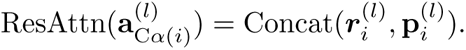

After obtaining representations for each atom, we apply a mean pooling layer over representations of all alpha carbons **a**_C*α*(*i*)_ to get protein representations **h**. An anti-symmetric prediction head is then applied to guarantee that back mutations would have the exact opposite predicted ΔΔ*G*_bind_ values:

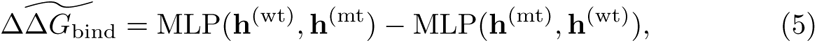

where **h**^(wt)^ and **h**^(mt)^ denote the representations for wild type and mutant complexes and 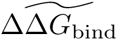 is the predicted ΔΔ*G*_bind_ from our GearBind model.

### Modeling energy landscape of proteins via noise contrastive estimation

As paired binding free energy change data is of relatively small size, it would be beneficial to pretrain GearBind with massive protein structural data. The high-level idea of our pretraining method is to model the distribution of native protein structures, which helps identify harmful mutations yielding unnatural structures. Denoting a protein structure as **x**, its distribution can be modeled with Boltzmann distribution as:

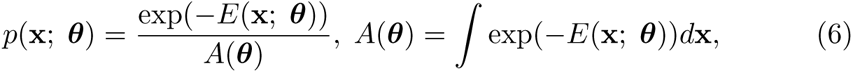

where ***θ*** denotes learnable parameters in our encoder, *E*(**x**; ***θ***) denotes the energy function for the protein *x* and *A*(***θ***) is the partition function to normalize the distribution. The energy function is predicted by applying a linear layer on the GearBind representations **h**(**x**) of protein **x**:

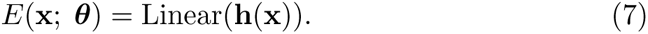

Given the observed dataset {**x**_1_,…, **x***_T_*} from PDB, our objective is to maximize the probability of these samples:

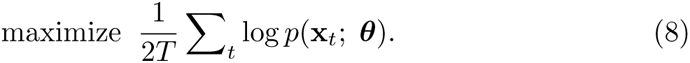

However, direct optimization of this objective is intractable, since calculating the partition function requires integration over the whole protein structure space. To address this issue, we adopt a popular method for learning energy-based models called noise contrastive estimation [24]. For each observed structure **x***_t_*, we sample a negative structure **y***_t_* and then the problem can be transformed to a binary classification task, *i.e.*, whether a sample is observed in the dataset or not.

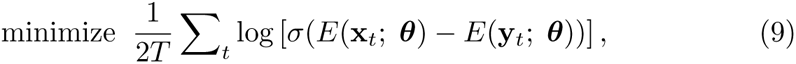

where *σ*(*·*) denotes the sigmoid function for calculating the probability for a sample **x***_t_* belonging to the positive class. We could see that the above training objective tries to push down the energy of the positive examples (*i.e.* the observed structures) while pushing up the energy of the negative samples (*i.e.* the mutant structures.

For negative sampling, we perform random single-point mutations on the corresponding positive samples and then generate its conformation by keeping the backbone unchanged and sampling side-chain torsional angles at the mutation site from a backbone-dependent rotamer library [25]. Besides, to further enhance the model’s capability to distinguish structural noises, we randomly choose 30% residues to randomly rotate torsional angles when generating negative samples.

After pretraining on the CATH database, we finetune the GearBind encoder on downstream tasks for prediction to avoid overfitting.

### Cross Validation on SKEMPI

During cross validation, a model is trained and tested five times, each time using a different subset as the test set and the remaining four subsets as the training set. Results are calculated for each test set, and their mean and standard error of mean are reported as the final cross validation performance. During the process of cross-validation, each individual data point is incorporated into the test set precisely once. This ensures that a comprehensive “test result table” is compiled, which includes predictive values for each data point when it is part of the test set. Subsequent performance analysis are done by splitting this table by various criteria and evaluate performance on each subset. After cross validation on SKEMPI, we obtain five sets of model parameters. During inference, we use the mean of the predicted values of these five checkpoints as the model prediction result.

### Baseline implementation details

#### FoldX

In this work, we use FoldX 4 [9] for mutant structure generation. First, each PDB file is processed with the RepairPDB command for structural corrections. Then, the wild-type, mutant structure pair is built using the BuildModel command. We use the AnalyseComplex command to get the FoldX ΔΔ*G*_bind_prediction based on the wild-type and mutant structures.

#### Flex-ddG

We run Flex-ddG with its official implementation at https://github.com/Kortemme-Lab/flex_ddG_tutorial. Each PDB file is processed with PDBFixer v1.8 [37]. Using the default parameters, we sample 35 structure models for each mutation, with the number of backrub trails set to 35000 and the energy function set to fa talaris2014. The final ΔΔ*G*_bind_ values are predicted with a generalized additive model that reweights the score terms.

#### GearBind(+P)

We implement GearBind with the TorchDrug library [41]. For message passing, we employed a 4-layer GearBind model with a hidden dimension of 128. Regarding edge message passing, the connections between edges are categorized into 8 bins according to the angles between them. To predict the ΔΔ*G*_bind_ value from graph representations, we utilized a 2-layer MLP.

The model was trained using the Adam optimizer with a learning rate of 1e-4 and a batch size of 8. The training process is performed on 1 A100 GPU for 40 epochs. For pretraining, we use the same architecture with 4-layer GearBind model with a hidden dimension of 128. The pretraining was conducted using the Adam optimizer with a learning rate of 5e-4 and a batch size of 8, employing 4 A100 GPUs for 10 epochs.

#### Bind-ddG

To ensure a fair comparison, we re-implement and re-train the Bind-ddG model on our SKEMPI splits. We following the configuration of the original implementation to set the dimensions of hidden and pair representations at 128 and 64, respectively. Our validation performance indicates that the optimal configuration for our setup includes a two-layer geometric attention mechanism and a four-layer MLP predictor. We trained the model using an Adam optimizer with a learning rate of 1e-4 and a batch size of 8, on a single A400 GPU, for a total of 40 epochs.

### In silico affinity maturation of CR3022 and anti-5T4 UdAb

PDB 6XC3 [42], in which chains H and L comprise antibody CR3022 and chain C is the SARS-CoV-2 RBD, was chosen as the starting complex for CR3022 affinity maturation. To better simulate the CR3022 interaction with Omicron RBD, we constructed the complex structures for BA.4 and BA.1.1 mutants with SWISS-MODEL [35]. We then performed saturation mutagenesis on the CDRs of CR3022 and generated mutant structures using FoldX [9] and Flex-ddG [10]. Specifically, residues 26-35, 50-66, 99-108 from the heavy chain H and residues 24-40, 56-62, 95-103 from the light chain L are mutated. This totals 1400 single-point mutations (if we count the self-mutations). We use our ensemble model to rank the mutations and select the top-ranked mutants for synthesis and subsequent experimental validation. Mutations are ranked by the modified *z*-score (where values are subtracted by the median rather than the mean to be less sensitive to outliers) averaged across multiple ΔΔ*G*_bind_ prediction methods.

An unpublished complex structure was used to optimize anti-5T4 UdAb. As the single-domain antibody binding two distinct epiopes on 5T4 (Fig. 5b), anti-5T4 UdAb has a larger interface region compared to traditional antibodies. After analyzing its interface with 5T4, we decided to perform saturation mutagenesis on residues 1,3,25,27-30,31-33,39-45,52-57,59,91-93,95,99,100-102,103,105,110,112,115-117. This totals 780 single-point mutations (if we count the self-mutations) that goes through the same ranking and selection strategies as described above.

### Antigen preparation

The gene encoding SARS-CoV RBD was synthesized by Genscript (Nanjing, China) and subcloned into pSectag 2B vector with C-terminal human IgG1 Fc fragment and AviTag. The recombinant vector was transfected to Expi 293 cells and cultured at 37*^◦^*C for 5 days, followed by centrifugation at 2, 200 *× g* for 20 minutes. The supernatant was harvested and filtered through a 0.22 µm vacuum filter. The protein G resin (Genscript) was loaded into the column, washed by PBS, and flow the supernatant through to fully combine the resin. Then the targeted protein was eluted with 0.1 M glycine (pH 3.0) and neutralized with 1 M Tris-HCL (pH 9.0), followed by buffer-exchanged and concentrated with phosphate buffered saline (PBS) using an Amicon ultra centrifugal concentrator (Millipore) with a molecular weight cut-off of 3 kDa. Protein concentration was measured using the NanoDrop 2000 spectrophotometer (Thermo Fisher), and protein purity was examined by sodium dodecyl sulfate-polyacrylamide gel electrophoresis (SDS-PAGE). The Delta RBD protein was purchased from Vazyme (Nanjing, China) and Omicron S protein was purchased from ACROBiosystems (Beijing, China). The biotinylated human TPBG / 5T4 and human TPBG/5T4-Fc antigen was purchased from ACROBiosystems (Beijing, China).

### Preparation for mutant and wild-type CR3022 antibodies

The heavy chain and light chain genes of different CR3022 antibodies were synthesized and subcloned into expression vector pcDNA 3.4 in IgG1 format. These constructed vectors were transfected into CHO cells and purified by Protein A. All antibodies were produced by Biointron Biological Inc. (Shanghai, China).

### Generation of mutant and wild-type anti-5T4 UdAbs

The pComb3x vector encoding the gene of wild-type anti-5T4 UdAb was constructed in previous work and preserved in our laboratory. All anti-5T4 UdAb mutants with single-point mutation were constructed with QuickMutation™ Site-directed Gene Mutagenesis Kit (Beyotime, Shanghai, China) following the manufacturer’s protocol. The expression of different mutant and wild-type anti-5T4 UdAbs were performed in *E. coli* HB2151 bacterial culture at 30*^◦^*C for 16 h accompanied by 1 mM isopropyl b-D-1-thiogalactopyranoside (IPTG). The cells were harvested and lysed by polymyxin B at 30*^◦^*C for 0.5 h, followed by centrifugation at 8, 800 *× g* for 10 minutes. The supernatant was collected, filtered through 0.8 µm polyethersulphone membranes by sterile syringes and purified by Ni-NTA (Smart Lifesciences) following the manufacturer’s instructions. Briefly, the filtered supernatant was loaded over the column with Ni-NTA. The resin was washed by washing buffer (10 mM Na2HPO4, 10 mM NaH2PO4 [pH 7.4], 500 mM NaCl, and 20 mM imidazole), and proteins were eluted in elution buffer (10 mM Na2HPO4, 10 mM NaH2PO4 [pH 7.4], 500 mM NaCl, and 250 mM imidazole). The collected pure fractions were immediately buffer exchanged into PBS and concentrated with Amicon ultra centrifugal concentrators (Millipore). Protein concentration was measured using the NanoDrop 2000 spectrophotometer (Thermo Fisher), and protein purity was examined by SDS-PAGE.

### Enzyme-linked immunosorbent assay (ELISA)

For comparison of different CR3022 mutants, the RBD of Delta (B.1.617.2) strain and S protein of Omicron (B.1.1.529) strain at 100 ng per well was coated in 96 wells half area microplates (Corning #3690) overnight at 4 *^◦^*C. The antigen coated plate was washed by three times with PBST (PBS with 0.05% Tween-20) and blocked with 3% MPBS (PBS with 3% skim milk) at 37 *^◦^*C for 1 h. Following three times washing with PBST, 50 µL of three-fold serially diluted antibody in 1% MPBS was added and incubated at 37 *^◦^*C for 1.5 h. The HRP-conjugated anti-Fab and anti-Fc (Sigma-Aldrich) secondary antibodies were used for the detection of different tested antibodies. After washing with PBST for 5 times, the enzyme activity was measured after the addition of ABTS substrate (Invitrogen) for 15 min. The data was acquired by measuring the absorbance at 405 nm using a Microplate Spectrophotometer (Biotek) and the EC_50_ (concentration for 50% of maximal effect) was calculated by GraphPad Prism8.0 software. To verify different UdAb mutants, the same experimental protocol as mentioned above was adopted. Briefly, the human TPBG/5T4-Fc antigen was coated in 96 wells half area microplates, then blocked with 3% MPBS and added serial diluted antibodies. The HRP-conjugated anti-Flag (Sigma-Aldrich) secondary antibody was used, followed by adding ABTS substrate and detected at 405 nm. The reported EC_50_ value is the mean value from three duplicates on a single experiment.

### Bio-layer Interferometry (BLI) binding assay

The binding kinetics of different antibodies to SARS-CoV-2 Omicron S and 5T4 antigens were measured by BLI on an Octet-RED96 (ForteBio). Briefly, the his-tagged Omicron S protein at 8 µg/ml and biotinylated 5T4 protein at 5 µg/ml were separately loaded onto Ni-NTA and streptavidin-coated (SA) biosensors. The antigen immobilized sensors were incubated with three-fold serially diluted CR3022 candidates or two-fold serially diluted anti-5T4 UdAbs starting at 300 nM in 0.02% PBST for 300 s for association, and then immersed into 0.02% PBST for another 300 s at 37 *^◦^*C for dissociation. All the curves were fitted by 1:1 binding model using the Data Analysis software 11.1. All *K*_D_ values were determined with *R*^2^ values of greater than 98% confidence level.

### Protein structure and ΔΔ*G*_bind_ contribution analysis

Protein structure analysis is conducted by python scripts. The antibody-antigen complex structure after mutation was obtained from Rosetta Flex-ddG relaxation [10]. The relaxed protein structure can provide more accurate side-chain conformations, which are critical for accurate contact and conformational analysis. The improved accuracy of such analyses enables a deeper understanding of the underlying binding mechanisms and can facilitate the identification of key characteristics involved in protein-protein interactions. The contribution scores are derived by using Integrated Gradients (IG) [43], a model-agnostic attribution method, on GearBind to obtain residue-level interpretation following [22]. All protein structure figures are created with PyMOL v3.0.

### Molecular dynamics simulation

For antibody mutation structural analysis, we conducted molecular dynamics simulations of the wild type and mutant antibody-antigen complex. Initial structures were taken from the Rosetta Flex-ddG relaxed structures used by GearBind. The LEaP module in the AMBER 22 suite was used for building starting structures and adding ions and solvent for the simulation [44]. The protonation states of the molecules were kept at the default settings as assigned by LEaP during the initial structure preparation. All systems were simulated with the ff19SB protein force field [45] and solvated in boxes of water with the OPC3 [46] solvent model. Simulated systems were solvated using a 10 Angstrom solvent box. All bonds involving hydrogen atoms were constrained with the SHAKE algorithm [47]. The particle mesh Ewald (PME) algorithm was used to calculate long-range electrostatic interactions [48]. Initial structures were relaxed with maximum 10,000 steps of minimization before convergence, then subjected to heating for 20 ps and equilibrating for 10 ps in the NPT ensemble with PMEMD. The CUDA version of PMEMD was used to accelerate the simulations [49]. The simulation temperature was set to room temperature 298K. All systems were simulated for 1 µs production MD with one replica, and samples in the first 50 ns were not used for analysis. CPP-TRAJ module in AMBERTools were used to analysis the simulation results, including calculating root mean square deviation (RMSD), room mean square fluctuation (RMSF), hydrogen bonding, and dihedral angle [50].

## Data Availability

The raw SKEMPI database can be accessed via https://life.bsc.es/pid/skempi2. The CATH database can be accessed via https://www.cathdb.info/. The raw HER2 binders data can be accessed via https://github.com/AbSciBio/unlocking-de-novo-antibody-design/blob/main/spr-controls.csv.

## Code Availability

The GearBind inference code, the trained model checkpoints and the dataset preprocessing scripts are available via https://github.com/DeepGraphLearning/GearBind under the Apache 2.0 License. They can also be accessed via Zenodo [51].

## Author contributions

J.T. conceptualized the study and supervised the project. T.Y. and Y.W. co-supervised the project. H.C. investigated related work, co-led model development, processed the datasets and *in silico* results, and led manuscript writing. Z.Z. investigated related work, led model development. B.Z. led the analysis of the structures and MD trajectories of the proposed mutants. M.W. was in charge of the *in vitro* experiments and results analysis with the help of Q.L. and Y.Z. All participated in manuscript writing.

