## Supplementary materials for "Pretrainable Geometric Graph Neural Network for Antibody Affinity Maturation"

#### Supplementary Text

#### Detailed analysis on SEKMPI

We plotted the performance of all benchmarked models against the number of mutations (Fig. S2), wild-type and mutant residue types (Fig. S3-S5), test data difficulty (Fig. 2a, b), wild-type & mutant binding (Fig. S7, S8) and absolute ΔΔG values (Fig. S9).

**Number of mutations.** Though one would expect that multi-mutations are more difficult to model than single-point mutations, Fig. S2 shows that the performance of all benchmarked models does not degrade significantly as the number of mutations in the protein complex goes up, indicating their capability of handling multi-mutations.

**Wild-type and mutant residue types.** Fig. S3a illustrates the data distribution for each (wild-type AA, mutant AA) pair in the SKEMPI dataset, revealing a significant bias towards alanine scanning data, where residues are predominantly mutated to alanine (A). The dataset shows that no other (wild-type AA, mutant AA) pair accumulates more than 50 observations, leading to less reliable performance metrics for these pairs as shown in Fig. S4, S5. Additionally, Fig. S3b highlights that mutations to tyrosine (Y) and leucine (L) frequently result in highly positive ΔΔG values, underscoring the critical role these residues play in binding affinity.

**Test data difficulty.** Fig. 2a, b uses boxplots to visualize the performance of various models on data points with various difficulties. PDB codes in SKEMPI are categorized into "easy" (50+ similar data points in training set), "medium" (1-50), and "hard" (0) targets based on the number of training data points having a high structural similarity (TM-score > 0.8) to it. Deep learning models, namely Bind-ddG, GearBind and GearBind+P, enjoys superior performance compared to physics-based methods such as FoldX and Flex-ddG on easy targets, but the table turns when we move to the hard targets, showing room for improvement in generalization capabilities. We also note that pretraining improves GearBind's performance, especially on easy targets. The ensemble method, overall, achieves the best results.

**Wild-type and mutant binding.** In Figure S6, we plot a heatmap visualizing the entry count of SKEMPI subsets binned by wild-type & mutant binding. The most frequent binding level changes are "medium → medium", "weak → weak", "medium → weak" and "strong → strong". Fig. S7 and S8 report the performance on SKEMPI subsets with different wild-type & mutant binding levels. We note that deep learning models perform significantly better than FoldX and Flex-ddG for weak-to-strong and medium-to-strong mutations, though this could possibly relate to these bins having relatively small sizes (Fig. S6) or the corresponding data have close relatives in the training dataset.

**Absolute ΔΔG values.** Fig. S9 plots the performance on SKEMPI subsets with low, medium and high |ΔΔG| values. All models perform better when the binding level changes more drastically. GearBind achieves outstanding performance in this region, with a PearsonR value of 0.707, compared to FoldX's 0.411, showing its potential of identifying mutations that could significantly increase the binding affinity. When the |ΔΔG| is small, predictions from all methods have very low correlation with experimental ΔΔG values. This could be due to the noises in data or a deficiency of current tools in modeling weaker interactions.

### ESM performance discussion

Protein language models (PLMs) are powerful pretrained models that have great potential in protein function prediction and optimization. Table S3 shows the spearman correlation between the values predicted by the ESM models (zero-shot prediction via likelihood ratio) and the experimental ddG. As we can see from the table, the SpearmanR scores are fairly high (about 0.9) within the ESM models, but all models have negative Spearman correlation to the experimental ddG. This indicates that, though zero-shot prediction of protein language models can be used to enhance affinity and stability in some cases, it is not a generally reliable method to rank the binding affinity of protein complexes. See Fig. S12 for per-PDB performance comparison between the ESM models, GearBind and the ensemble model.

During this study, we also tried fine-tuning ESM on SKEMPI. However, when we apply this model on the CR3022 affinity maturation task, we find it systematically overweighs mutations to W and Y (we are aware that W and Y are enriched in antibody paratopes[1], but blindly suggesting mutation to these residues at all positions is clearly incorrect), perhaps due to bias in the training data (Fig. S3). We chose not to use this fine-tuned ESM model in our affinity maturation pipeline. This shows that sequence-based language models, with only amino acid types provided as input, could be less robust to spurious correlations in data.

[1] Reis, Pedro BPS, et al. "Antibody-antigen binding interface analysis in the big data era." *Frontiers in Molecular Biosciences* 9 (2022): 945808.

### Inference time cost comparison

We benchmarked the inference time cost of each model in Table S11. FoldX and Flex-ddG requires seconds and hours, respectively, to compute the ddG calculation for a single mutant. In contrast, deep learning models such as GearBind, GearBind+P, and Bind-ddG complete their inference in a fraction of a second.

It is important to note that the deep learning models rely on the FoldX-modelled mutant structure for their inference. Therefore, ensembling them with FoldX ddG predictions adds only a negligible time cost to the overall inference process while resulting in significant performance improvements.

### Evaluation on another independent test set

We collected 9 BLI-measured ddG values for single-point mutation on the Spike RBD from the paper[1], and evaluated the PearsonR and SpearmanR of various models on this small test set (Table S12). GearBind and GearBind+P gave the best performance among all benchmarked models.

However, from the MAE and RMSE values, we could see that none of the benchmarked models were accurate in terms of the predicted $\Delta\Delta G$ value. In Table S13, we listed 9 more mutations with no precise $\Delta\Delta G$ figures as the mutant binding is too weak to be measured. The $\Delta\Delta G$ values of these mutations are expected to be greater than 2, but no model predicted any ddG value greater than 2. This case study showcases the unsolved challenges for existing $\Delta\Delta G_{\text{bind}}$ prediction methods.

[1] Muecksch, Frauke, et al. "Affinity maturation of SARS-CoV-2 neutralizing antibodies confers potency, breadth, and resilience to viral escape mutations." Immunity 54.8 (2021): 1853-1868.

### Equilibration of the simulated system

To analyze the equilibration of the simulated system, we plotted the RMSD of two simulated systems. As shown in Fig. S18 and S19, the RMSD reached a steady state after 50 ns of simulation (the first 50 ns of simulation were discarded in further structure analysis). Additionally, we presented the overall number of hydrogen bonds over simulation time in Fig. S29 and S30. The numbers of overall hydrogen bonds also reached steady states within 50 ns of simulation.

### Supplementary Tables

**Table S1.** Cross validation performance of different methods on the single-point mutation subset of SKEMPI ($n$ = 4068). For each metric, we report the mean and standard error of the mean. "+P" means "with geometric pretraining".

| | **MAE** | **RMSE** | **PearsonR** | **SpearmanR** |
|---|---|---|---|---|
| **FoldX** | 1.124 ± 0.087 | 1.759 ± 0.130 | 0.432 ± 0.018 | <u>0.435 ± 0.009</u> |
| **Flex-ddG** | 1.054 ± 0.084 | 1.620 ± 0.130 | 0.454 ± 0.032 | 0.421 ± 0.022 |
| **Bind-ddG** | 1.065 ± 0.061 | 1.506 ± 0.076 | 0.547 ± 0.054 | 0.371 ± 0.037 |

| | | | | |
|---|---|---|---|---|
| **GearBind** | <u>0.946 ± 0.055</u> | <u>1.384 ± 0.057</u> | <u>0.650 ± 0.038</u> | 0.422 ± 0.030 |
| **GearBind+P** | **0.917 ± 0.045** | **1.361 ± 0.041** | **0.668 ± 0.042** | **0.466 ± 0.035** |
| **Ensemble** | 0.874 ± 0.061 | 1.293 ± 0.073 | 0.713 ± 0.024 | 0.565 ± 0.026 |

**Table S2.** Ablation study of GearBind, quantified by cross validation performance on SKEMPI (*n* = 5729). For each metric, we report the mean and standard error of the mean.

| | PearsonR | SpearmanR | MAE | RMSE |
|---|---|---|---|---|
| **FoldX** | 0.491 ± 0.007 | 0.526 ± 0.011 | 1.364 ± 0.134 | 2.027 ± 0.170 |
| **Flex-ddG** | 0.497 ± 0.034 | 0.484 ± 0.020 | 1.236 ± 0.101 | 1.849 ± 0.150 |
| **Bind-ddG** | 0.581 ± 0.037 | 0.443 ± 0.041 | 1.255 ± 0.096 | 1.759 ± 0.125 |
| **GearBind** | 0.659 ± 0.030 | 0.498 ± 0.033 | 1.143 ± 0.088 | 1.639 ± 0.103 |
| **GearBind+P** | 0.676 ± 0.041 | 0.525 ± 0.046 | 1.115 ± 0.072 | 1.611 ± 0.075 |
| **Ensemble** | 0.729 ± 0.016 | 0.643 ± 0.030 | 1.028 ± 0.080 | 1.503 ± 0.101 |
| **GearBind w/o resi MP** | 0.627 ± 0.054 | 0.482 ± 0.045 | 1.176 ± 0.058 | 1.683 ± 0.055 |
| **GearBind w/o edge MP** | 0.631 ± 0.029 | 0.431 ± 0.031 | 1.221 ± 0.095 | 1.751 ± 0.115 |
| **GearBind w/o side-chain atoms** | 0.599 ± 0.035 | 0.421 ± 0.030 | 1.271 ± 0.099 | 1.821 ± 0.122 |
| **GearBind w/o relational graph** | 0.519 ± 0.055 | 0.384 ± 0.033 | 1.305 ± 0.115 | 1.882 ± 0.159 |
| **RGCN (w/ relational graph)** | 0.573 ± 0.049 | 0.454 ± 0.050 | 1.262 ± 0.055 | 1.765 ± 0.064 |

**Table S3.** Pairwise Spearman correlations between ESM predictions and experimental $\Delta\Delta G$ values of the SKEMPI dataset (*n* = 5729).

| | esm-1b | esm-1v-m1 | esm-1v-m2 | esm-1v-m3 | esm-1v-m4 | esm-1v-m5 | $\Delta\Delta G$ |
|---|---|---|---|---|---|---|---|
| **esm-1b** | 1 | 0.893 | 0.897 | 0.878 | 0.861 | 0.894 | -0.188 |
| **esm-1v-m1** | 0.893 | 1 | 0.922 | 0.917 | 0.904 | 0.924 | -0.186 |
| **esm-1v-m2** | 0.897 | 0.922 | 1 | 0.909 | 0.913 | 0.937 | -0.215 |
| **esm-1v-m3** | 0.878 | 0.917 | 0.909 | 1 | 0.916 | 0.916 | -0.196 |
| **esm-1v-m4** | 0.861 | 0.904 | 0.913 | 0.916 | 1 | 0.919 | -0.207 |
| **esm-1v-m5** | 0.894 | 0.924 | 0.937 | 0.916 | 0.919 | 1 | -0.205 |
| **ddG** | -0.188 | -0.186 | -0.215 | -0.196 | -0.207 | -0.205 | 1 |

**Table S4.** PearsonR between model predictions on SKEMPI (*n* = 5729).

| | FoldX | Flex-ddG | Bind-ddG | GearBind | GearBind+P | Ensemble |
|---|---|---|---|---|---|---|
| **FoldX** | 1.000 | 0.587 | 0.402 | 0.392 | 0.371 | 0.749 |
| **Flex-ddG** | 0.587 | 1.000 | 0.416 | 0.434 | 0.429 | 0.717 |
| **Bind-ddG** | 0.402 | 0.416 | 1.000 | 0.727 | 0.731 | 0.814 |
| **GearBind** | 0.392 | 0.434 | 0.727 | 1.000 | 0.893 | 0.816 |

| | | | | | | |
|---|---|---|---|---|---|---|
| **GearBind+P** | 0.371 | 0.429 | 0.731 | 0.893 | 1.000 | 0.811 |
| **Ensemble** | 0.749 | 0.717 | 0.814 | 0.816 | 0.811 | 1.000 |

**Table S5.** SpeamanR between model predictions on SKEMPI ($n$ = 5729).

| | **FoldX** | **Flex-ddG** | **Bind-ddG** | **GearBind** | **GearBind+P** | **Ensemble** |
|---|---|---|---|---|---|---|
| **FoldX** | 1.000 | 0.639 | 0.389 | 0.423 | 0.419 | 0.771 |
| **Flex-ddG** | 0.639 | 1.000 | 0.400 | 0.449 | 0.438 | 0.732 |
| **Bind-ddG** | 0.389 | 0.400 | 1.000 | 0.550 | 0.557 | 0.743 |
| **GearBind** | 0.423 | 0.449 | 0.550 | 1.000 | 0.732 | 0.692 |
| **GearBind+P** | 0.419 | 0.438 | 0.557 | 0.732 | 1.000 | 0.697 |
| **Ensemble** | 0.771 | 0.732 | 0.743 | 0.692 | 0.697 | 1.000 |

**Table S6.** SKEMPI data filtered out due to high variance (std > 1) between measurements of the same mutation. The largest difference between measurements is 5.3 kcal/mol, which would surely confuse the model if used in training.

| **pdb_chains** | **mutations** | **count** | **mean** | **std** | **min** | **25%** | **50%** | **75%** | **max** |
|---|---|---|---|---|---|---|---|---|---|
| **1A22_A_B** | **IB64A** | 3 | 1.350 | 1.055 | 0.132 | 1.036 | 1.940 | 1.959 | 1.977 |
| | **RB11A** | 3 | 1.523 | 1.079 | 0.278 | 1.197 | 2.115 | 2.146 | 2.177 |
| **1AO7_ABC_DE** | **DD26W,RD27F,GD28T,SD51M,SD94T** | 2 | -2.014 | 1.032 | -2.743 | -2.379 | -2.014 | -1.649 | -1.284 |
| | **EA166A** | 4 | 2.199 | 1.322 | 0.877 | 1.307 | 2.027 | 2.919 | 3.865 |
| **1BP3_A_B** | **EA170A** | 2 | 1.831 | 2.348 | 0.170 | 1.001 | 1.831 | 2.661 | 3.491 |
| | **HA18A** | 2 | 1.549 | 1.926 | 0.187 | 0.868 | 1.549 | 2.230 | 2.911 |
| | **HA21A** | 2 | 1.246 | 2.014 | -0.178 | 0.534 | 1.246 | 1.958 | 2.671 |
| | **RA163A** | 2 | 3.023 | 1.277 | 2.121 | 2.572 | 3.023 | 3.475 | 3.926 |
| **1BRS_A_D** | **DD39A** | 2 | 6.790 | 1.213 | 5.932 | 6.361 | 6.790 | 7.218 | 7.647 |
| **1CHO_EFG_I** | **TI14E** | 2 | 5.582 | 1.067 | 4.827 | 5.205 | 5.582 | 5.959 | 6.336 |
| **1DAN_HL_UT** | **VU113A** | 2 | 0.690 | 1.243 | -0.189 | 0.251 | 0.690 | 1.130 | 1.569 |
| **1FC2_C_D** | **IC27A** | 2 | 3.729 | 2.185 | 2.184 | 2.957 | 3.729 | 4.502 | 5.274 |
| **1JTG_A_B** | **EA79K** | 2 | 3.239 | 1.406 | 2.245 | 2.742 | 3.239 | 3.736 | 4.233 |
| | **EA85A** | 2 | 1.377 | 3.794 | -1.305 | 0.036 | 1.377 | 2.719 | 4.060 |
| **2WPT_A_B** | **SB67A** | 2 | -0.134 | 1.011 | -0.849 | -0.491 | -0.134 | 0.223 | 0.581 |
| **3MZG_A_B** | **HA167A,HB188A** | 9 | 2.653 | 1.278 | -0.237 | 2.801 | 3.329 | 3.391 | 3.493 |
| **3S9D_A_B** | **RA26A** | 3 | 5.123 | 1.322 | 3.680 | 4.548 | 5.415 | 5.845 | 6.275 |
| **4NKQ_C_AB** | **DB176A** | 3 | 1.511 | 1.016 | 0.422 | 1.049 | 1.676 | 2.055 | 2.434 |

**Table S7.** Validation performance of GearBind with and without pre-training for retrieving wild-type proteins.

| Validation performance | HITS@1 ↑ | HITS@5 ↑ | HITS@10 ↑ | Mean Rank ↓ | Mean Reciprocal Rank ↑ |
|---|---|---|---|---|---|
| **GearBind (random init)** | 0.000 | 0.1813 | 0.4313 | 11.9625 | 0.1294 |
| **GearBind (pre-trained)** | **0.5750** | **0.6000** | **0.7312** | **5.9812** | **0.6186** |

**Table S8.** RBD sequences of tested SARS-CoV-2 mutants used in this research. Residues on the interface to CR3022 (i.e. having a heavy atom within 5 Angstroms of CR3022) are highlighted in orange.

```
                    334   340       350       360       370       380       390
wt                  NLCPFGEVFNATRFASVYAWNRKRISNCVADYSVLYNSASFSTFKCYGVSPTKLNDLCFTNVYADS
                    ||||||||||||||||||||||||||||||||||||||||||||||||||||||||||||||||||
delta (B.1.617.2)   NLCPFGEVFNATRFASVYAWNRKRISNCVADYSVLYNSASFSTFKCYGVSPTKLNDLCFTNVYADS
                    |||||.||||||.||||||||||||||||||||||||.|.|.||||||||||||||||||||||||
omicron (BA.1.1)    NLCPFDEVFNATKFASVYAWNRKRISNCVADYSVLYNLAPFFTFKCYGVSPTKLNDLCFTNVYADS
                    ||||||||||||.||||||||||||||||||||||||.||||.|||||||||||||||||||||||
omicron (BA.4)      NLCPFDEVFNATRFASVYAWNRKRISNCVADYSVLYNFAPFFAFKCYGVSPTKLNDLCFTNVYADS

                    400       410       420       430       440       450       460
wt                  FVIRGDEVRQIAPGQTGKIADYNYKLPDDFTGCVIAWNSNNLDSKVGGNYNYLYRLFRKSNLKPFE
                    |||||||||||||||||||||||||||||||||||||||||||||||||||.||||||||||||||
delta (B.1.617.2)   FVIRGDEVRQIAPGQTGKIADYNYKLPDDFTGCVIAWNSNNLDSKVGGNYNYRYRLFRKSNLKPFE
                    |||||||||||||||||.||||||||||||||||||||||.|||||.||||.||||||||||||||
omicron (BA.1.1)    FVIRGDEVRQIAPGQTGNIADYNYKLPDDFTGCVIAWNSNKLDSKVSGNYNYLYRLFRKSNLKPFE
                    |||||.||.|||||||||||||||||||||||||||||||||||||.||||.||||||||||||||
omicron (BA.4)      FVIRGNEVSQIAPGQTGNIADYNYKLPDDFTGCVIAWNSNKLDSKVGGNYNYRYRLFRKSNLKPFE

                        470       480       490       500       510       520
wt                  RDISTEIYQAGSTPCNGVEGFNCYFPLQSYGFQPTNGVGYQPYRVVVLSFELLHAPATVCGPK
                    ||||||||||||.||||||||||||||||||||||||||||||||||||||||||||||||||
delta (B.1.617.2)   RDISTEIYQAGSKPCNGVEGFNCYFPLQSYGFQPTNGVGYQPYRVVVLSFELLHAPATVCGPK
                    |||||||||||.||||||.||||||||.||.|.||.|||.||||||||||||||||||||||
omicron (BA.1.1)    RDISTEIYQAGNKPCNGVAGFNCYFPLRSYSFRPTYGVGHQPYRVVVLSFELLHAPATVCGPK
                    ||||||||||||||||||||.||||||.||.||||||||||||||||||||||||||||||||
omicron (BA.4)      RDISTEIYQAGNKPCNGVAGVNCYFPLQSYGFRPTYGVGHQPYRVVVLSFELLHAPATVCGPK
```

**Table S9.** Similarity of the RBD structure used for CR3022 affinity maturation (pdb 6XC3_C) to all SKEMPI chains, measured by TMalign.

| ref_pdb | query_pdb | len1 | len2 | len_aln | rmsd | seq_id | tm_score1 | tm_score2 |
|---|---|---|---|---|---|---|---|---|
| 6xc3_C.pdb | 2AJF_E.pdb | 193 | 174 | 172 | 1.3 | 0.692 | 0.851 | 0.94 |
| 6xc3_C.pdb | 4ZS6_A.pdb | 193 | 207 | 173 | 3.26 | 0.179 | 0.716 | 0.674 |
| 6xc3_C.pdb | 1FSS_A.pdb | 193 | 532 | 126 | 5.69 | 0.071 | 0.365 | 0.172 |
| 6xc3_C.pdb | 1MAH_A.pdb | 193 | 533 | 119 | 5.22 | 0.059 | 0.357 | 0.167 |
| 6xc3_C.pdb | 2NYY_A.pdb | 193 | 1267 | 117 | 5.58 | 0.051 | 0.346 | 0.077 |
| 6xc3_C.pdb | 1B41_A.pdb | 193 | 531 | 121 | 5.61 | 0.091 | 0.346 | 0.165 |
| 6xc3_C.pdb | 2NZ9_A.pdb | 193 | 1267 | 117 | 5.57 | 0.06 | 0.345 | 0.077 |
| 6xc3_C.pdb | 4NM8_E.pdb | 193 | 317 | 104 | 4.9 | 0.048 | 0.339 | 0.233 |

| | | | | | | | | |
|---|---|---|---|---|---|---|---|---|
| 6xc3_C.pdb | 4FZA_B.pdb | 193 | 281 | 114 | 5.58 | 0.079 | 0.338 | 0.259 |
| 6xc3_C.pdb | 3NVN_A.pdb | 193 | 383 | 121 | 5.85 | 0.066 | 0.337 | 0.208 |

**Table S10.** Similarity of the RBD structure used for 5T4 affinity maturation to all SKEMPI chains, measured by TMalign.

| ref_pdb | query_pdb | len1 | len2 | len_aln | rmsd | seq_id | tm_score1 | tm_score2 |
|---|---|---|---|---|---|---|---|---|
| 5T4_A.pdb | 4Y61_B.pdb | 283 | 235 | 234 | 2.63 | 0.261 | 0.72946 | 0.86338 |
| 5T4_A.pdb | 4YEB_B.pdb | 283 | 321 | 243 | 3.2 | 0.235 | 0.72815 | 0.65039 |
| 5T4_A.pdb | 1Z7X_W.pdb | 283 | 460 | 257 | 3.91 | 0.16 | 0.70976 | 0.46596 |
| 5T4_A.pdb | 1A4Y_A.pdb | 283 | 460 | 254 | 3.86 | 0.154 | 0.70469 | 0.46184 |
| 5T4_A.pdb | 3M63_A.pdb | 283 | 953 | 207 | 5.74 | 0.077 | 0.46253 | 0.17373 |
| 5T4_A.pdb | 4O27_A.pdb | 283 | 325 | 203 | 5.63 | 0.099 | 0.45296 | 0.40912 |
| 5T4_A.pdb | 4NZW_A.pdb | 283 | 309 | 191 | 5.37 | 0.063 | 0.44696 | 0.41805 |
| 5T4_A.pdb | 3M62_A.pdb | 283 | 955 | 184 | 5.11 | 0.065 | 0.43821 | 0.1601 |
| 5T4_A.pdb | 4FZA_A.pdb | 283 | 324 | 199 | 5.66 | 0.075 | 0.4364 | 0.39562 |
| 5T4_A.pdb | 1MAH_A.pdb | 283 | 533 | 200 | 6.1 | 0.085 | 0.42519 | 0.26357 |

**Table S11.** Inference time cost for each method, benchmarked on UdAb.

| | Task | Device | Inference time cost (s/mutation) |
|---|---|---|---|
| FoldX | Mutant structure modeling and ddG calculation with command "Pssm" | CPU | 7 |
| Flex-ddG | Mutant structure modeling and ddG calculation by running the full Flex-ddG pipeline | CPU | 3230 |
| GearBind / GearBind+P | ddG calculation with 5 models | GPU | 0.10 |
| Bind-ddG | ddG calculation with 5 models | GPU | 0.09 |

**Table S12.** Performance on single mutants in Muecksch et al. (n = 9).

| | PearsonR | SpearmanR | MAE | RMSE |
|---|---|---|---|---|
| FoldX | 0.02 | 0.17 | 1.95 | 2.48 |
| GearBind+P | 0.26 | 0.40 | 1.97 | 2.50 |
| GearBind | 0.40 | 0.63 | 1.94 | 2.45 |
| Bind-ddG | -0.72 | -0.40 | 2.26 | 3.06 |
| Flex-ddG | -0.08 | -0.02 | 1.98 | 2.65 |

**Table S13.** Ground truth $K_D$ values and $\Delta\Delta G$ model predictions of data points in Muecksch et al where the mutant antibodies are non-binding (n.b.).

| pdb_id | mutation | $K_D^{wt}$ (nM) | $K_D^{mt}$ (nM) | $\Delta\Delta G$ | FoldX | GearBind +P | GearBind | Bind-ddG | Flex-ddG |
|---|---|---|---|---|---|---|---|---|---|
| 7R8O | EB484K | 2.828 | n.b. | >2 | 1.300 | -0.369 | 0.020 | 0.033 | 1.018 |

| | | | | | | | | |
|---|---|---|---|---|---|---|---|---|
| 7R8O | KB417N,EB484K, NB501Y | 2.828 | n.b. | 1.281 | -0.109 | -0.025 | -0.186 | 1.463 |
| 7R8N_C052 | EA484K | 1.960 | n.b. | -1.403 | -0.298 | 0.124 | 0.438 | 0.281 |
| 7R8N_C052 | KA417N,EA484K, NA501Y | 1.960 | n.b. | -0.171 | -0.564 | -0.136 | 0.724 | 0.281 |
| 7R8N_C144 | EA484K | 33.150 | n.b. | -1.366 | -0.611 | 0.004 | 0.603 | -0.040 |
| 7R8N_C144 | KA417N,EA484K, NA501Y | 33.150 | n.b. | -0.479 | -0.621 | -0.067 | 0.537 | 0.489 |
| 7R8N_C144 | QA493R | 33.150 | n.b. | 0.130 | 0.140 | 0.219 | -1.565 | -0.592 |
| 7R8N | EA484K | 0.003 | n.b. | -1.302 | -0.453 | -0.163 | 0.445 | 0.273 |
| 7R8N | KA417N,EA484K, NA501Y | 0.003 | n.b. | -0.885 | -0.681 | -0.540 | 0.360 | 1.199 |

**Table S14.** Molecular dynamics simulation details

| System | Force field | Simulation time (us) | Number of trajectories | Number of water molecules | Ions |
|---|---|---|---|---|---|
| **CR3022 WT** | ff19SB+OPC3 | 1 | 1 | 46,172 | 7 Cl- |
| **CR3022 SH100D+ SH103Y+SL33R** | ff19SB+OPC3 | 1 | 1 | 46,133 | 7 Cl- |
| **UdAb WT** | ff19SB+OPC3 | 1 | 1 | 46,172 | 13 Na+ |
| **UdAb S57W** | ff19SB+OPC3 | 1 | 1 | 46,133 | 13 Na+ |

### Supplementary Figures

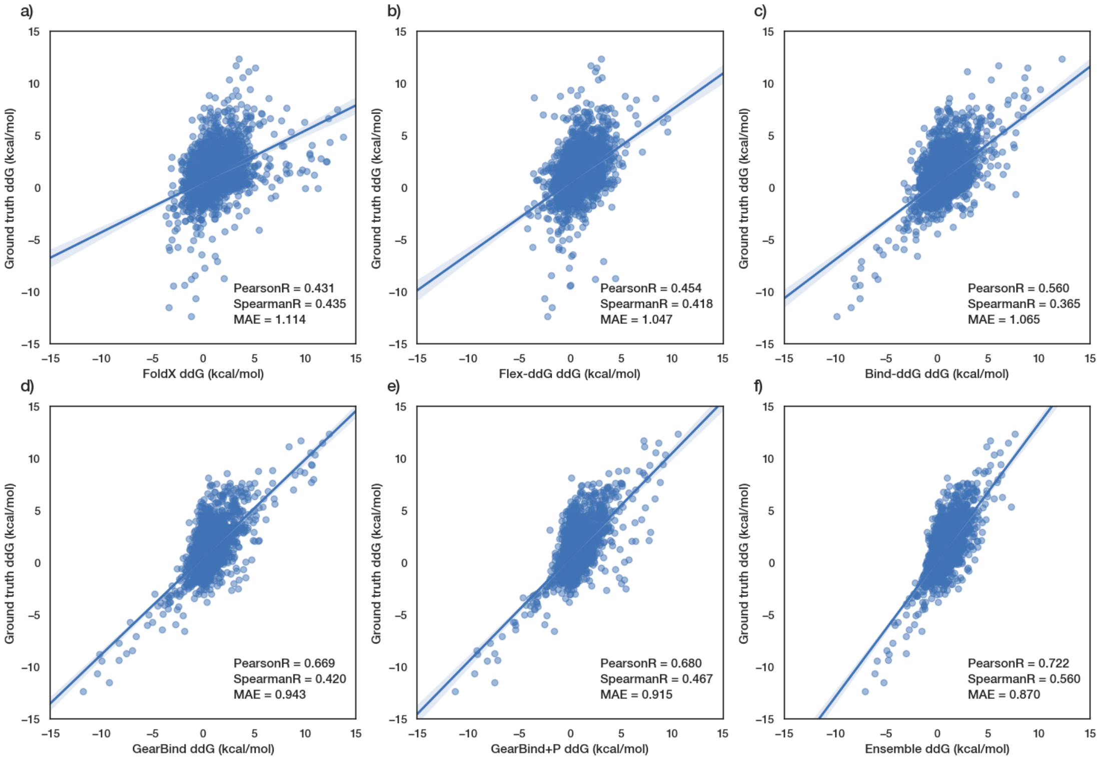

**Fig. S1.** Scatter plots of ground truth ΔΔ$G$'s versus ΔΔ$G$'s predicted by different models trained on SKEMPIv2 and tested on its single-mutant subset with split-by-complex cross validation ($n$ = 4068). Linear regression model fits are plotted with the data, with the light blue region around the line representing the 95% confidence interval. Pearson correlation, Spearman correlation and mean absolute error (MAE) values are annotated in each plot. The prediction of the ensemble model aligns best with the experimental data.

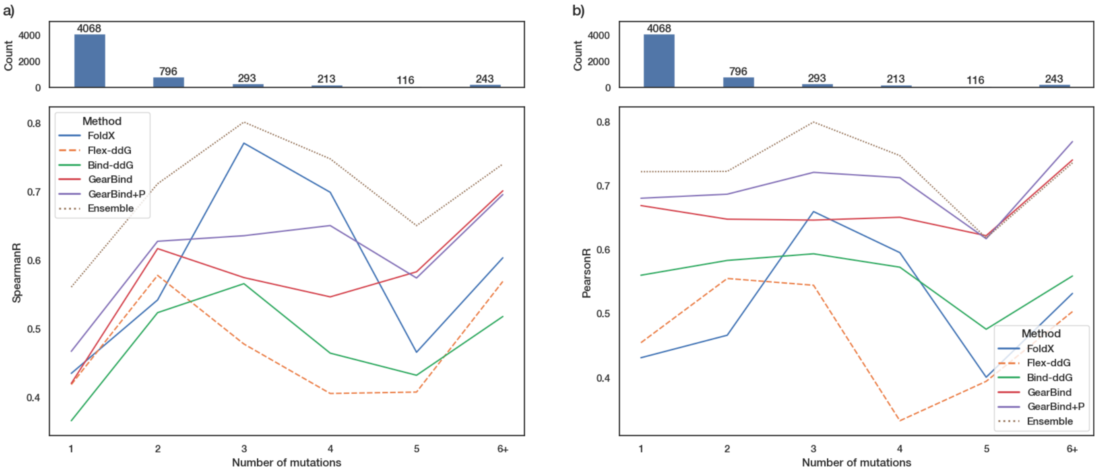


**Fig. S2.** Comparative analysis of model performance quantified by a) Spearman and b) Pearson correlation coefficients across SKEMPI subsets with varying mutation counts, illustrating the relationship between number of mutations in the complex and model accuracy. Models using FoldX-modelled structure are shown in straight lines; Flex-ddG which uses Rosetta modelled structure is shown in dashed lines; and the ensemble model is shown in dotted lines. Above each plot is a bar plot showing the number of SKEMPI data points carrying different numbers of mutations.

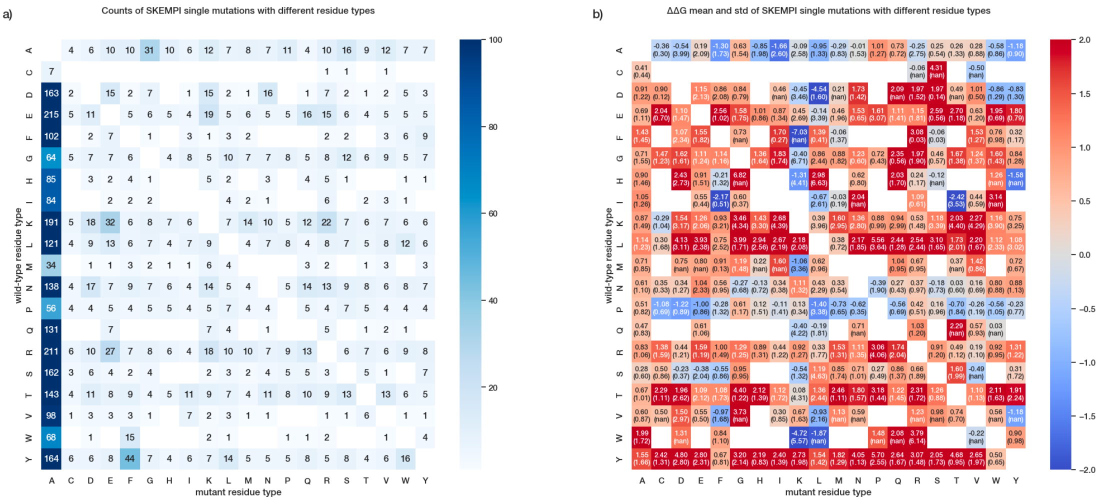


**Fig. S3.** Heatmap showing a) the number of observed mutations, b) mean and standard deviation (given in parentheses) of experimental ΔΔ$G$ for each (wild-type residue type, mutant residue type) pair in the SKEMPI single-mutant subset ($n$ =

4068). The non-observed residue type pairs are left blank.

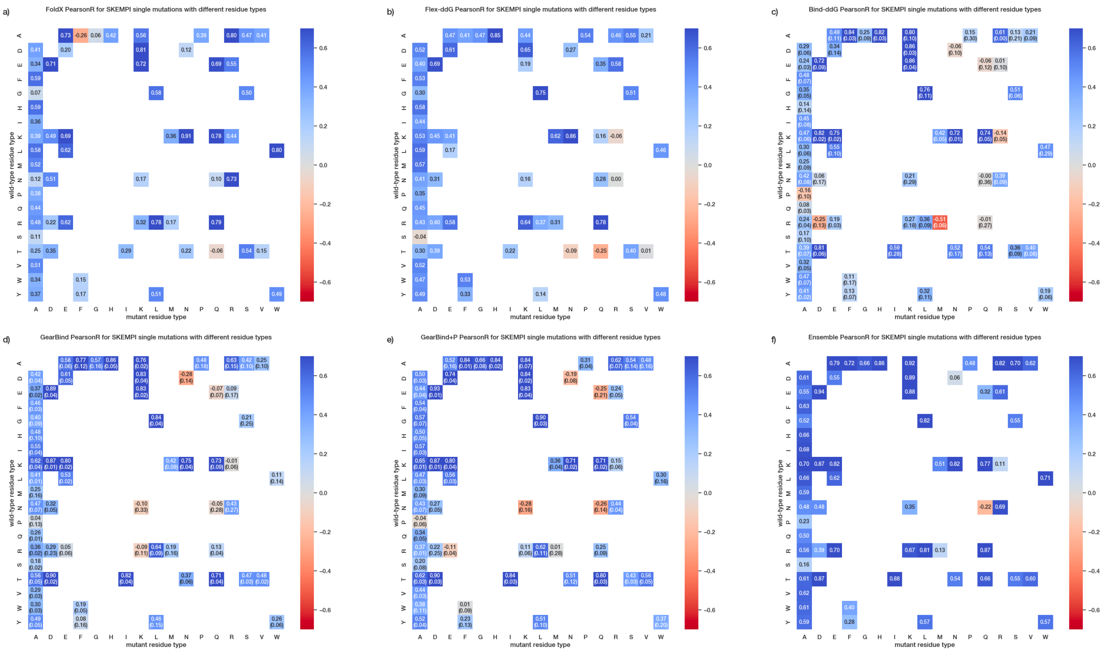


**Fig. S4.** Evaluation of model performance across (wild-type residue type, mutant residue type) pairs in the SKEMPI single-mutant subset (*n* = 2918; Residue type pairs with less than ten observations are omitted), visualized as PearsonR heatmaps. Each heatmap corresponds to a different predictive model, as indicated by the subfigure labels. a) FoldX, b) Flex-ddG, c) Bind-ddG, d) GearBind, e) GearBind+P, f) Ensemble model. For deep learning models (c-e), the heatmap is annotated with "mean (std)" where results are calculated from three training runs with different random seeds for weight initialization.

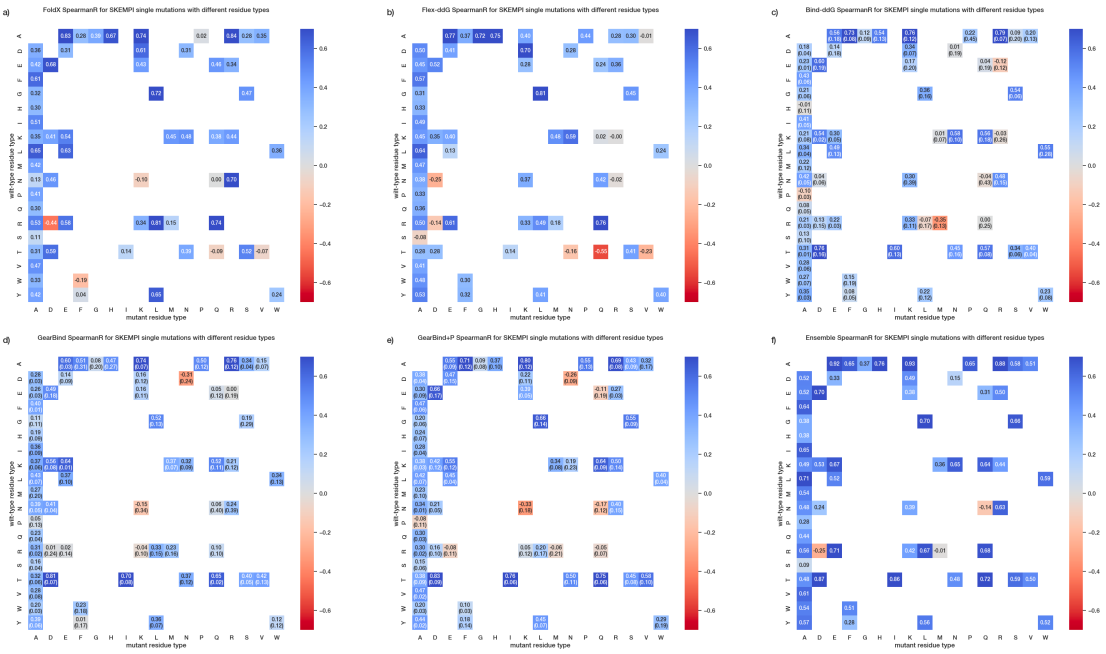

**Fig. S5.** Evaluation of model performance across (wild-type residue type, mutant residue type) pairs in the SKEMPI single-mutant subset ($n$ = 2918; Residue type pairs with less than ten observations are omitted), visualized as Spearman correlation coefficient (SpearmanR) heatmaps. Each heatmap corresponds to a different predictive model, as indicated by the subfigure labels. a) FoldX, b) Flex-ddG, c) Bind-ddG, d) GearBind, e) GearBind+P, f) Ensemble model. For deep learning models (c-e), the heatmap is annotated with "mean (std)" where results are calculated from three training runs with different random seeds for weight initialization.

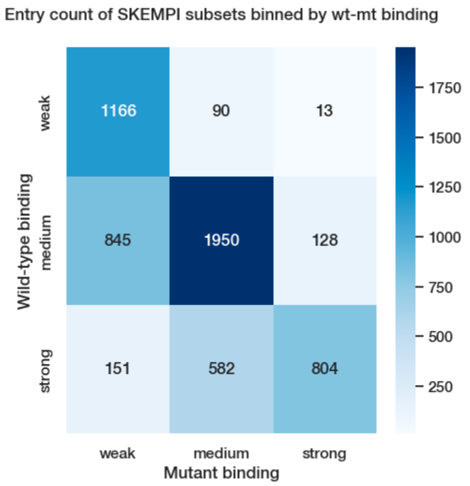


**Fig. S6.** Entry count of SKEMPI subsets ($n$ = 5729) binned by wt-mt binding. Binding affinities are binned into 3 levels: weak (>$10^{-7}$ M), medium ($10^{-10}$ to $10^{-7}$ M) and strong (<$10^{-10}$ M).

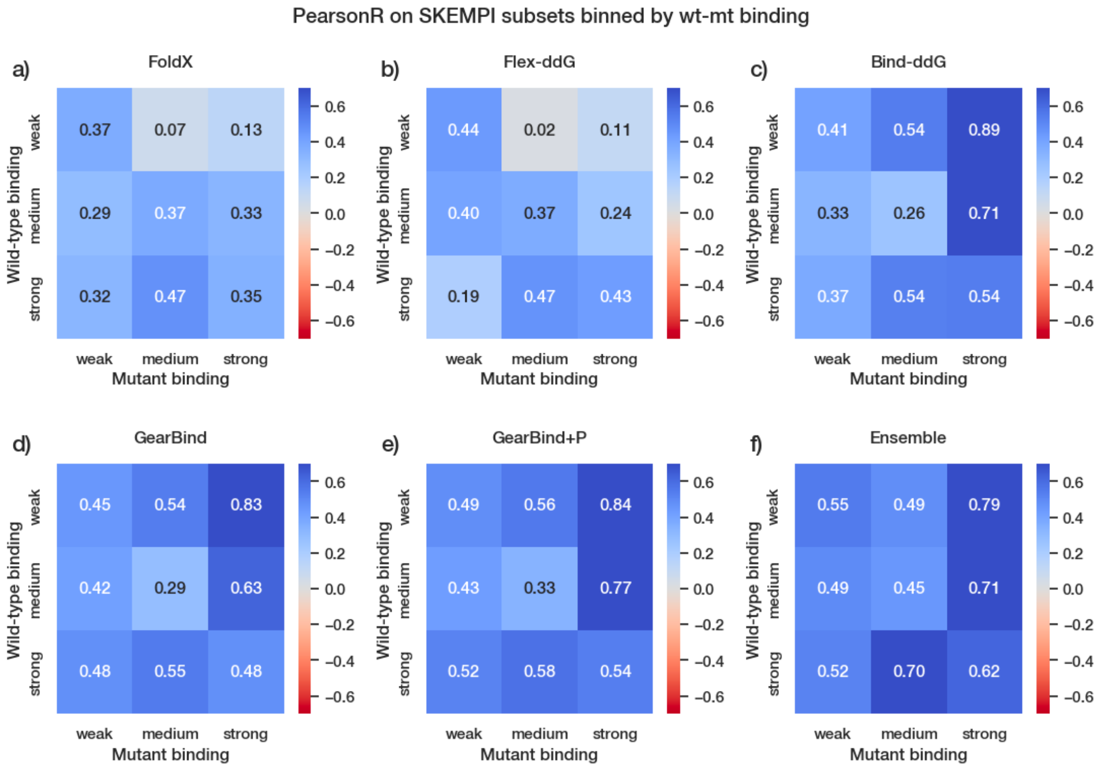


**Fig. S7.** Pearson correlation on SKEMPI data ($n$ = 5729) binned by wild-type and mutant binding level. The three levels are: weak (>$10^{-7}$ M), medium ($10^{-10}$ to $10^{-7}$ M) and strong (<$10^{-10}$ M).

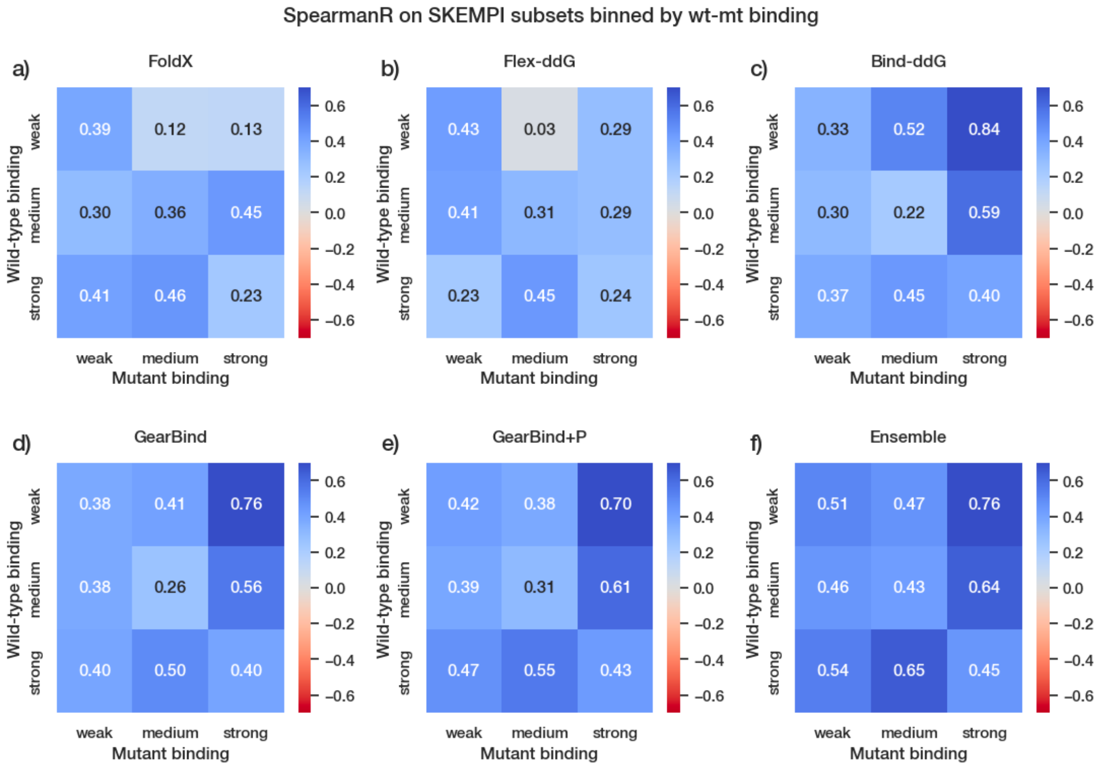


**Fig. S8.** Spearman correlation on SKEMPI data ($n$ = 5729) binned by wild-type and mutant binding level. The three levels are: weak (>$10^{-7}$ M), medium ($10^{-10}$ to $10^{-7}$ M) and strong (<$10^{-10}$ M).

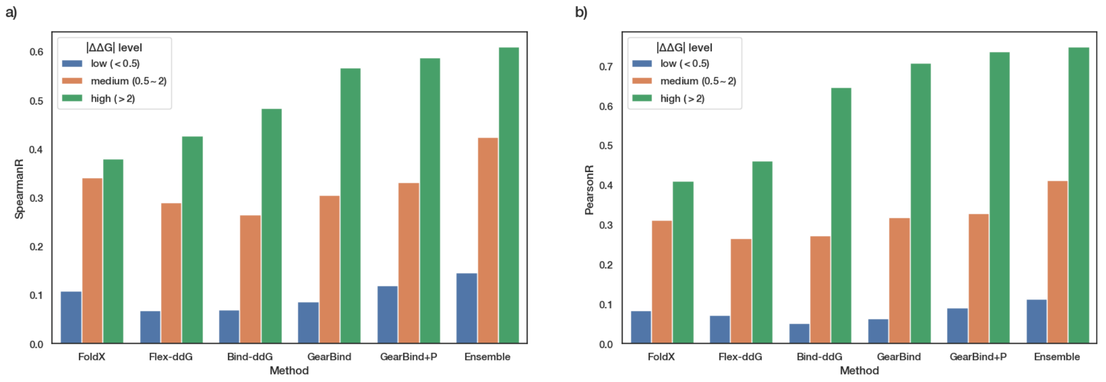


**Fig. S9.** Performance on SKEMPI data ($n$ = 5729) binned by the absolute $\Delta\Delta G$ value, as quantified by a) Spearman and b) Pearson correlations.

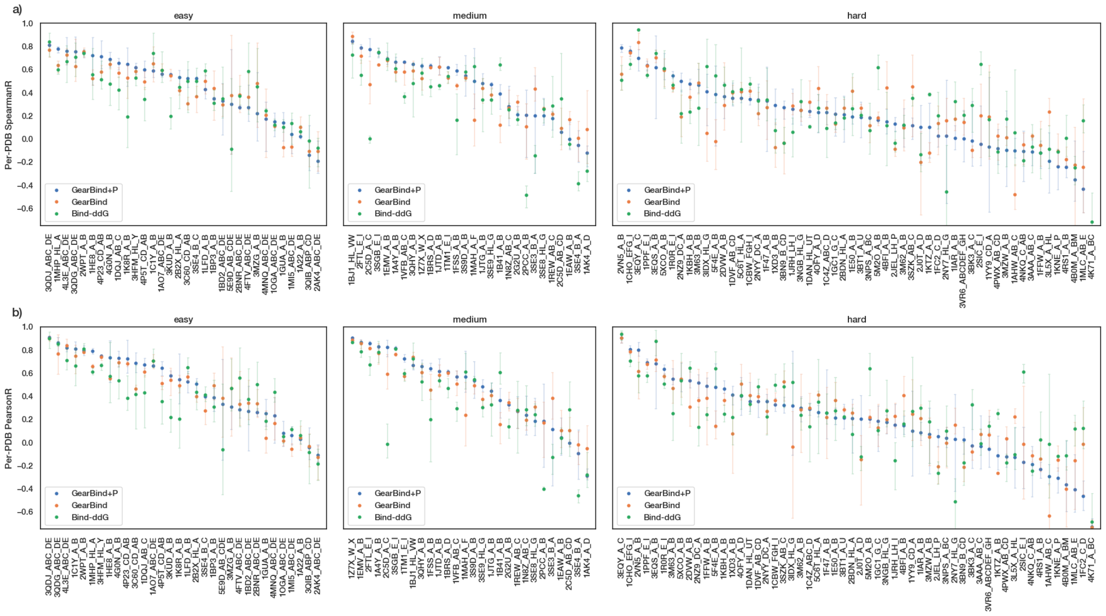


**Fig. S10.** Detailed analysis of Per-PDB a) Spearman and b) Pearson correlations between predictions of deep learning models and experimental data. PDBs with less than 10 experimental $\Delta\Delta G$ values are omitted. The models include GearBind+P (blue), GearBind (orange) and Bind-ddG (green). Error bars that show the 95% confidence interval are based on 3 training runs with different random seeds.

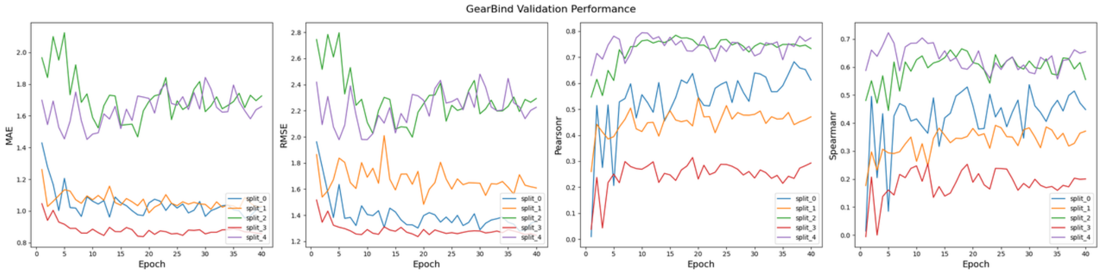


**Fig. S11.** Curves plotting four validation metrics (MAE, RMSE, PearsonR and SpearmanR) over training epochs during 5-fold cross-validation. Most of the validation metrics demonstrate improvement within the initial 10 epochs, followed by fluctuation throughout the subsequent 30 epochs. Consequently, extending training beyond 40 epochs appears unlikely to enhance performance significantly.

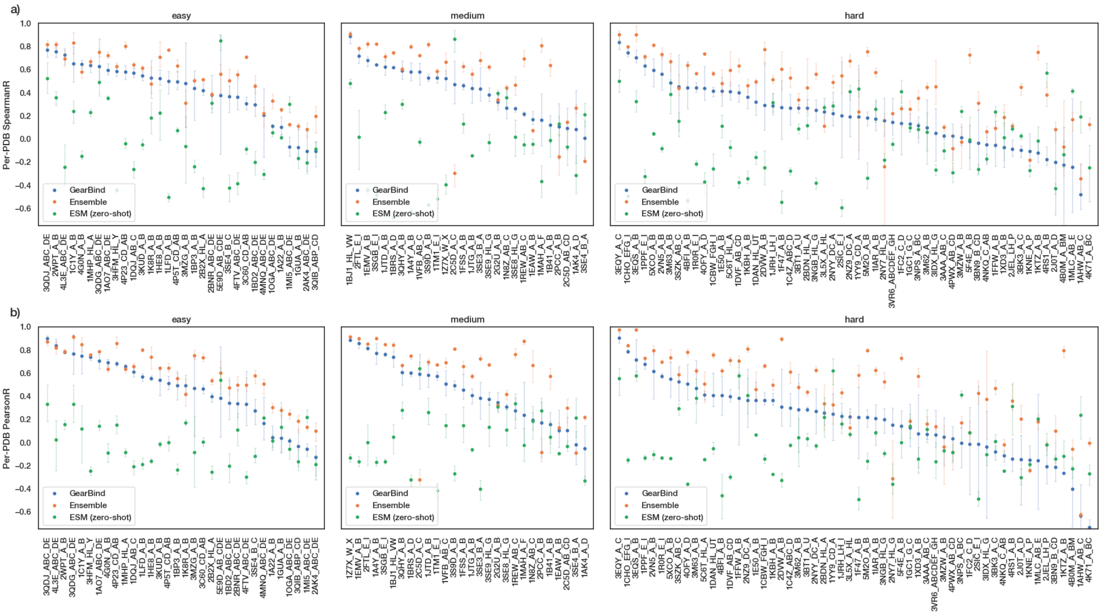


**Fig S12.** Evaluation of model robustness on the SKEMPI dataset via per-PDB a) Spearman and b) Pearson correlation coefficients between model predictions and experimental $\Delta\Delta G$ values. PDBs with less than 10 experimental $\Delta\Delta G$ values are omitted. Each model's mean correlation is denoted by a filled circle, and the 95% confidence intervals are depicted with error bars. ESM models (green) include ESM-1b (trained on UniRef50) and 5 versions of ESM-1v (trained on UniRef90).

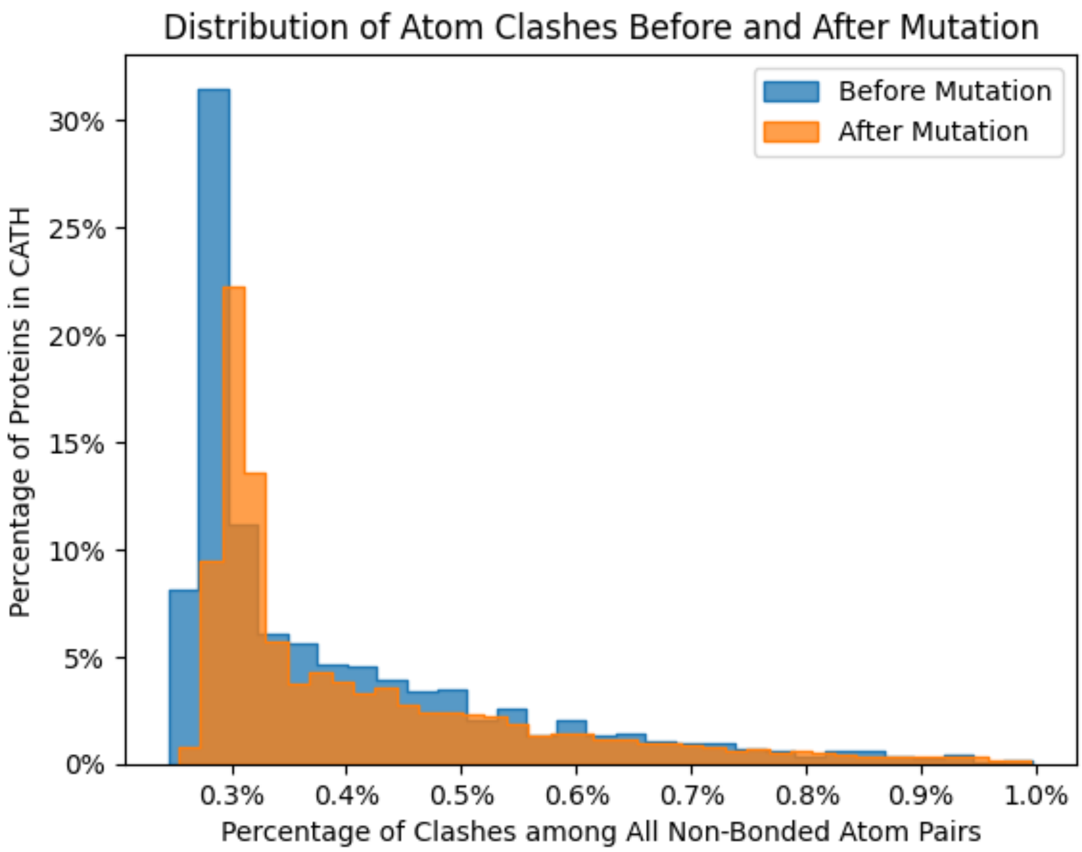


**Fig. S13.** Analysis of steric clashes introduced by the negative sampling scheme during pretraining. The numbers of atom clashes among all non-bonded atom pairs are calculated for both wild-type proteins and generated mutants. On average, there are 0.41% clashes of all pairs before mutation, which slightly increases to 0.44% after mutation.

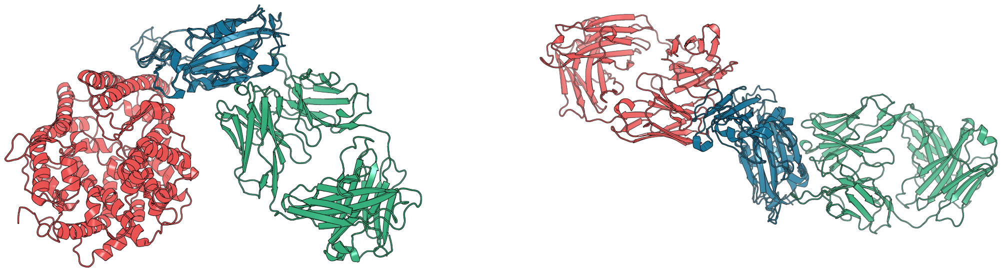

**Fig. S14.** Comparison of 6XC3 to similar proteins in SKEMPI. **Left**: PDB 2AJF, structure of the SARS RBD (blue) and the ACE2 (red) complex, aligned to PDB 6XC3, structure of the SARS-CoV-2 RBD (blue) and the CR3022 (green) complex. **Right**: PDB 4ZS6, structure of the MERS RBD (blue) and MERS-27 Fab (red) complex, aligned to PDB 6XC3. The RBDs are aligned using PyMOL CEAlign.

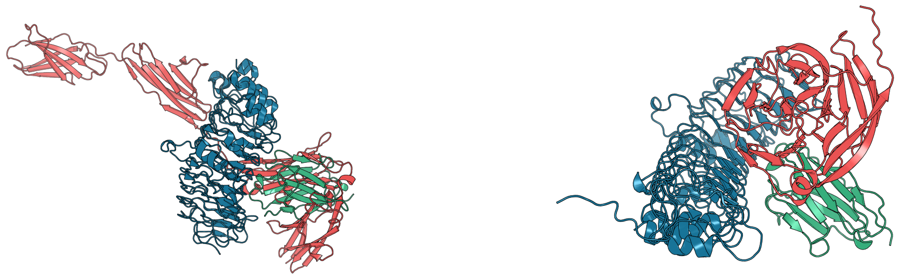

**Fig. S15.** Comparison of UdAb-5T4 to similar proteins in SKEMPI. **Left**: PDB 4Y61, structure of the Slitrk2 (blue) and the PTP delta (red) complex, aligned to the structure of the 5T4 (blue) and the UdAb (green) complex. **Right**: PDB 4YEB, structure of the FLRT3 (blue) and the Latrophilin-3 (red) complex, aligned to the structure of the 5T4 (blue) and the UdAb (green) complex. The 5T4-like domains are aligned using PyMOL CEAlign.

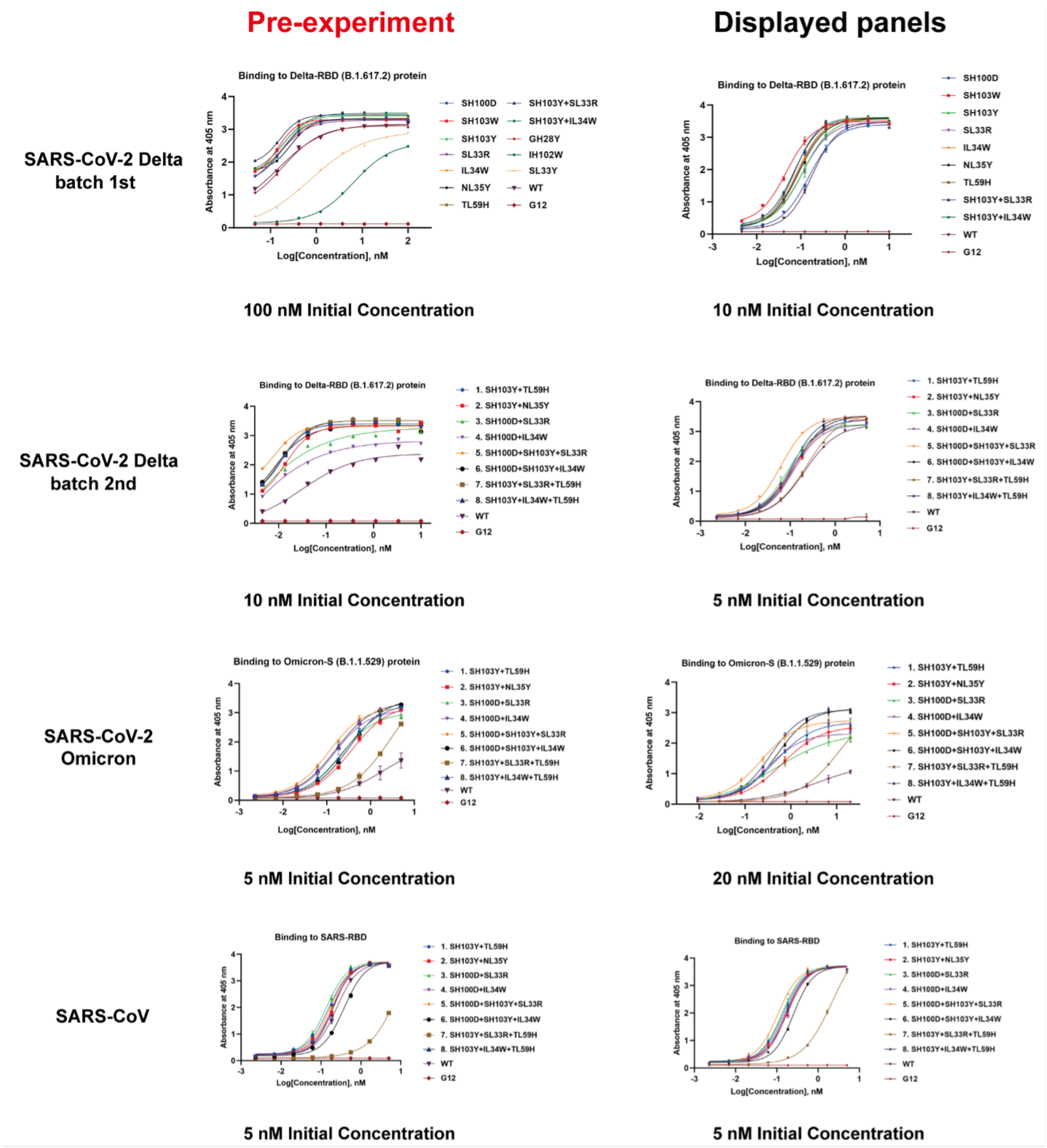


**Fig. S16. Left**: The pre-experiments conducted to determine the optimal starting concentration of tested antibodies. The initial concentration demonstrated in the display panel column is the determined optimal concentration. **Right**: The displayed panel in Fig. 3a, c, e, f corresponds to the pre-experiment in the left panel.

a)

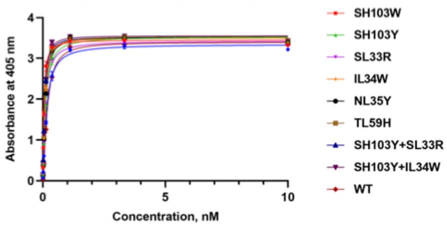


b)

| Mutation | EC50 (nM) | 95% Confidence Interval | Fold change |
|---|---|---|---|
| SH100D | 0.1453 | 0.1336~0.1581 | 1.19 |
| SH103W | 0.03987 | 0.03634~0.04375 | 3.22 |
| SH103Y | 0.1001 | 0.08882~0.1129 | 1.72 |
| SL33R | 0.09533 | 0.08739~0.1040 | 1.81 |
| IL34W | 0.07471 | 0.06944~0.08039 | 2.31 |
| NL35Y | 0.08302 | 0.07700~0.08952 | 2.08 |
| TL59H | 0.079 | 0.07315~0.08533 | 2.18 |
| SH103Y+SL33R | 0.06277 | 0.05767~0.06834 | 2.75 |
| SH103Y+IL34W | 0.06629 | 0.06141~0.07156 | 2.60 |
| CR3022-WT | 0.1725 | 0.1624~0.1833 | - |

c)

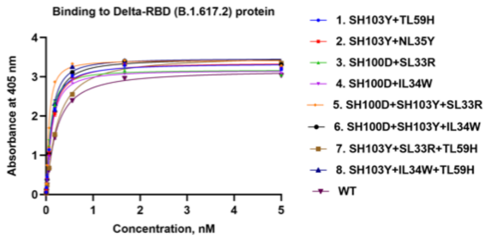


d)

| Mutation | EC50 (nM) | 95% Confidence Interval | Fold change |
|---|---|---|---|
| SH103Y+TL59H | 0.1048 | 0.09611~0.1145 | 1.97 |
| SH103Y+NL35Y | 0.1141 | 0.1040~0.1256 | 1.81 |
| SH100D+SL33R | 0.08496 | 0.07744~0.09336 | 2.44 |
| SH100D+IL34W | 0.1138 | 0.1058~0.1226 | 1.82 |
| SH100D+SH103Y+SL33R | 0.05736 | 0.05299~0.06213 | 3.61 |
| SH100D+SH103Y+IL34W | 0.1204 | 0.1110~0.1309 | 1.72 |
| SH103Y+SL33R+TL59H | 0.2292 | 0.2107~0.2504 | 0.90 |
| SH103Y+IL34W+TL59H | 0.1045 | 0.09621~0.1137 | 1.98 |
| CR3022-WT | 0.2069 | 0.1952~0.2196 | - |

e)

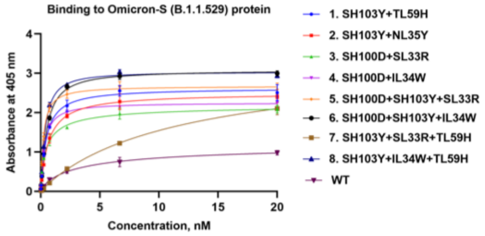


f)

| Mutation | EC50 (nM) | 95% Confidence Interval | Fold change |
|---|---|---|---|
| SH103Y+TL59H | 0.4464 | 0.3783~0.5337 | 7.90 |
| SH103Y+NL35Y | 0.648 | 0.5523~0.7744 | 5.44 |
| SH100D+SL33R | 0.4771 | 0.3716~0.6425 | 7.49 |
| SH100D+IL34W | 0.3072 | 0.2795~0.3389 | 11.47 |
| SH100D+SH103Y+SL33R | 0.217 | 0.1896~0.2497 | 16.24 |
| SH100D+SH103Y+IL34W | 0.4896 | 0.4524~0.5314 | 7.20 |
| SH103Y+SL33R+TL59H | 12.12 | 7.999~27.68 | 0.29 |
| SH103Y+IL34W+TL59H | 0.3521 | 0.3239~0.3835 | 10.01 |
| CR3022-WT | 3.525 | 2.181~7.898 | - |

g)

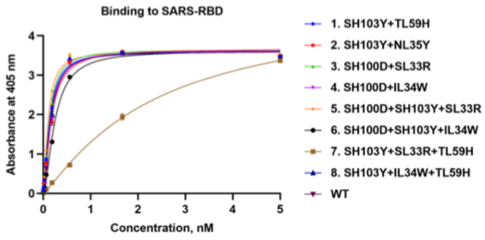


h)

| Mutation | EC50 (nM) | 95% Confidence Interval | Fold change |
|---|---|---|---|
| SH103Y+TL59H | 0.1313 | 0.1193~0.1446 | 1.22 |
| SH103Y+NL35Y | 0.1655 | 0.1471~0.1863 | 0.97 |
| SH100D+SL33R | 0.1252 | 0.1144~0.1371 | 1.28 |
| SH100D+IL34W | 0.1568 | 0.1412~0.1741 | 1.02 |
| SH100D+SH103Y+SL33R | 0.09874 | 0.09072~0.1075 | 1.63 |
| SH100D+SH103Y+IL34W | 0.2416 | 0.2227~0.2625 | 0.66 |
| SH103Y+SL33R+TL59H | 2.385 | 1.967~2.874 | 0.07 |
| SH103Y+IL34W+TL59H | 0.1465 | 0.1317~0.1630 | 1.10 |
| CR3022-WT | 0.1606 | 0.1471~0.1753 | - |

i)

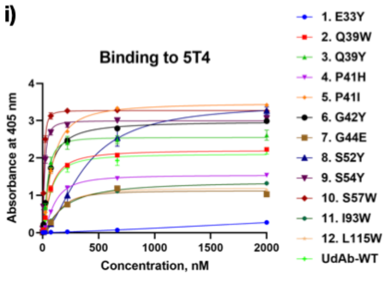


j)

| Mutation | EC50 (nM) | 95% Confidence Interval | Fold change |
|---|---|---|---|
| E33Y | No binding | - | - |
| Q39W | 70.67 | 66.04~75.80 | 0.99 |
| Q39Y | 42.37 | 38.25~46.98 | 1.65 |
| P41H | 104.3 | 99.83~109.2 | 0.67 |
| P41Y | 82.29 | 78.43~86.43 | 0.85 |
| G42Y | 58.1 | 53.86~62.86 | 1.20 |
| G44E | 139.1 | 119.9~161.0 | 0.50 |
| S52Y | 371.9 | 330.3~425.8 | 0.19 |
| S54Y | 17.18 | 16.06~18.38 | 4.07 |
| S57W | 12.6 | 12.15~13.06 | 5.55 |
| I93W | 190.3 | 177.4~205.1 | 0.37 |
| L115W | 147.1 | 134.3~161.3 | 0.48 |
| UdAb-WT | 69.94 | 60.25~82.49 | - |

**Fig. S17.** Hill equation fitted ELISA binding assay results for CR3022 and anti-5T4 UdAb mutants designed with a GearBind-based pipeline. On the left panels (a, c, e, g, i), the concentration-response curves alongside with $EC_{50}$ values evaluated from ELISA assays are displayed, with the center denoting the mean absorbance and error bars indicating the standard deviation from three technical duplicates. On the right panels (b, d, f, h, j), the fitted $EC_{50}$ values, their 95% confidence intervals and the fold changes in binding calculated as $EC_{50}^{(wt)}/EC_{50}^{(mt)}$ are displayed. Tested systems include the first-round CR3022 designs binding to Delta RBD (a, b); the second-round CR3022 designs binding to Delta RBD (c, d), Omicron S protein (e, f) and SARS-RBD (g, h); and single point mutants of anti-5T4 UdAb (i, j) binding to 5T4.

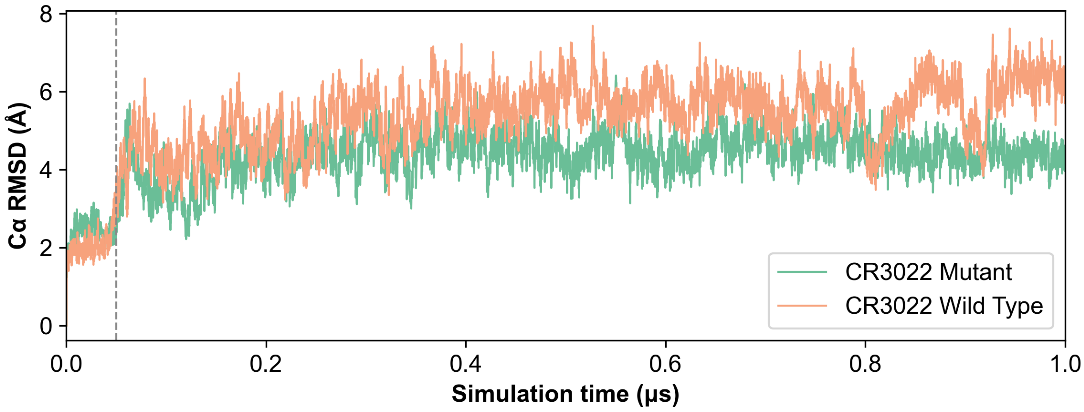


**Fig. S18.** Ca RMSD curve in the 1 μs MD simulation for the SARS-CoV-2 RBD-CR3022 complex. Only trajectories after 50 ns (marked by a vertical dashed line) are used in further analysis.

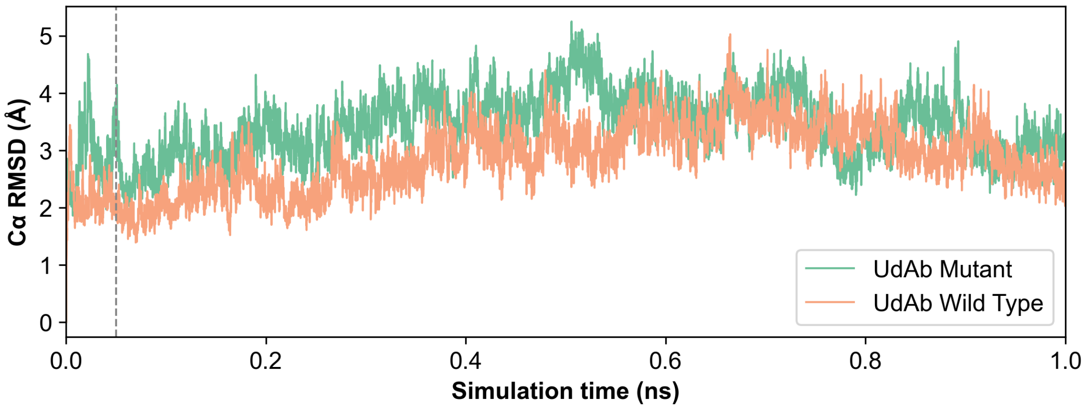


**Fig. S19.** Ca RMSD curve in the1 μs MD simulation for the 5T4-UdAb complex. Only trajectories after 50 ns (marked by a vertical dashed line) are used in further analysis.

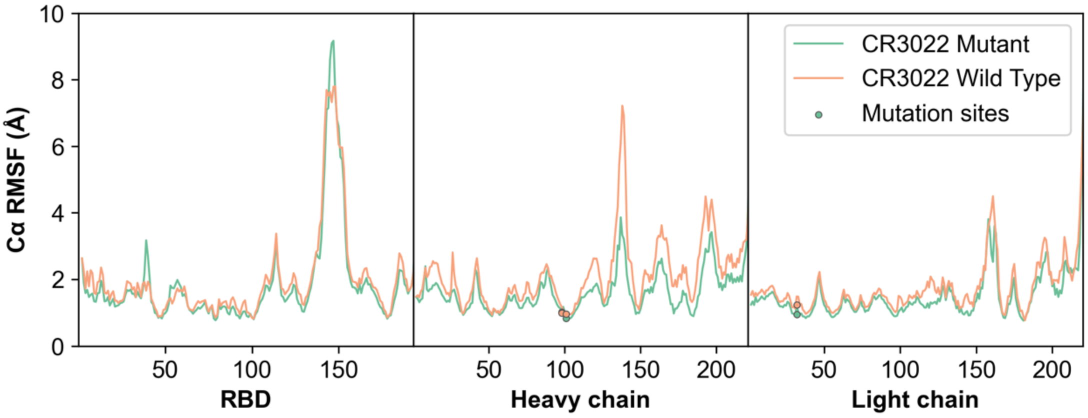


**Fig. S20.** Ca RMSF curve in the 1 μs MD simulation for the SARS-CoV-2 RBD-CR3022 complex, visualizing the fluctuation of the $C_{\alpha}$ atom in the simulation for wild type (orange) and SH100D, SH103Y, SL33R triple mutant (green). The mutation sites on the antibody chains are marked with dots.

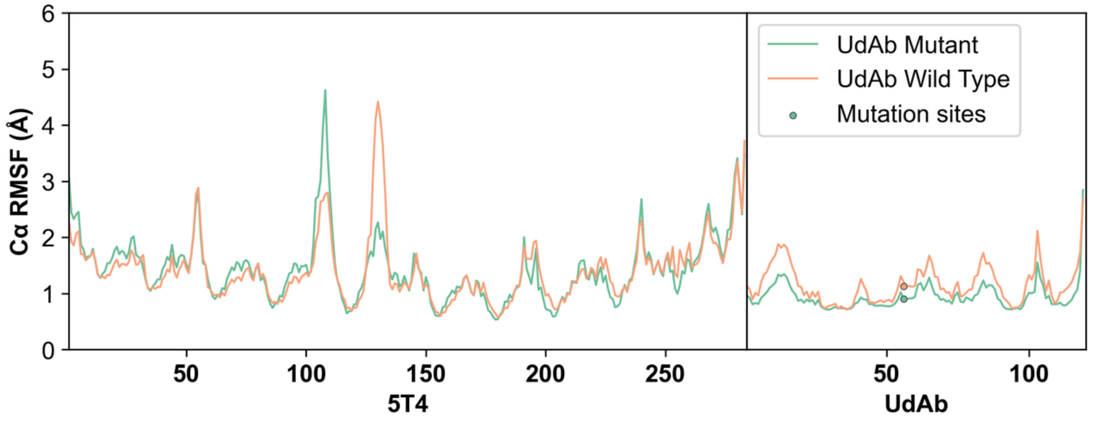


**Fig. S21.** Ca RMSF curve in the 1 µs MD simulation for the 5T4-UdAb complex, visualizing the fluctuation of the $C_\alpha$ atom in the simulation for wild type (orange) and the S57W mutant (green). The mutation sites on the antibody chain are marked with dots.

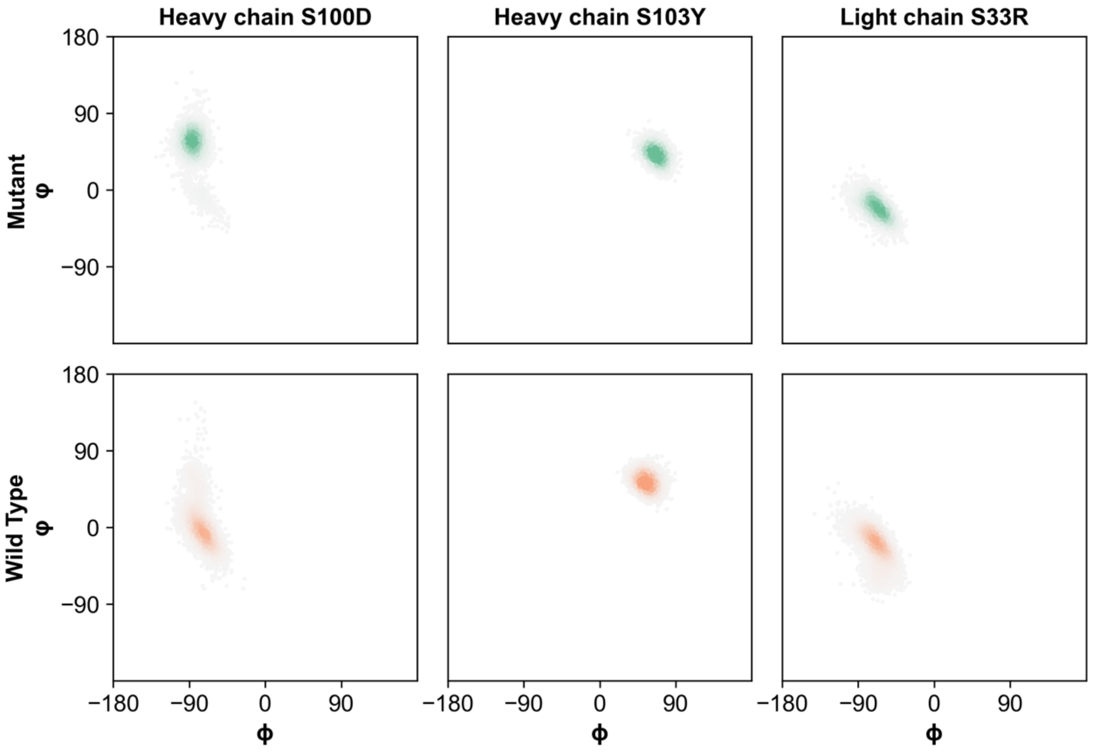


**Fig. S22.** Backbone torsion distribution of 3 residues at the CR3022 mutation sites after (top row) and before (bottom row) the SH100D+SH103Y+SL33R triple mutation. Density is fit by the gaussian kernel function, with high-density area colored by orange (wild type) and green (mutant), and low-density area colored by grey.

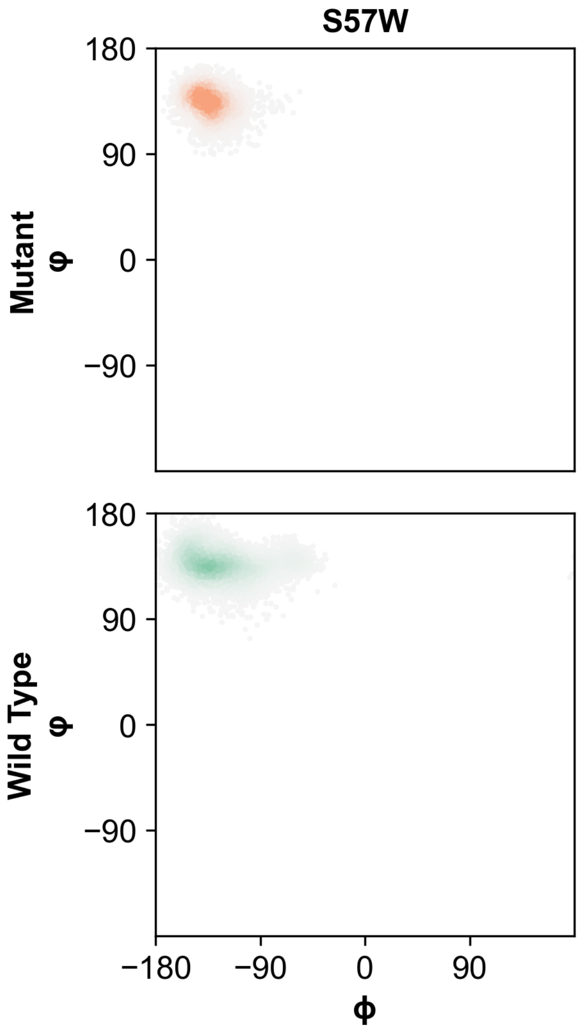


**Fig. S23.** Backbone torsion distribution of residue 57 in UdAb after (top row) and before (bottom row) the S57W mutation. Density is fit by gaussian kernel function, with high-density area colored by orange (mutant) and green (wild type), and low-density area colored by grey.

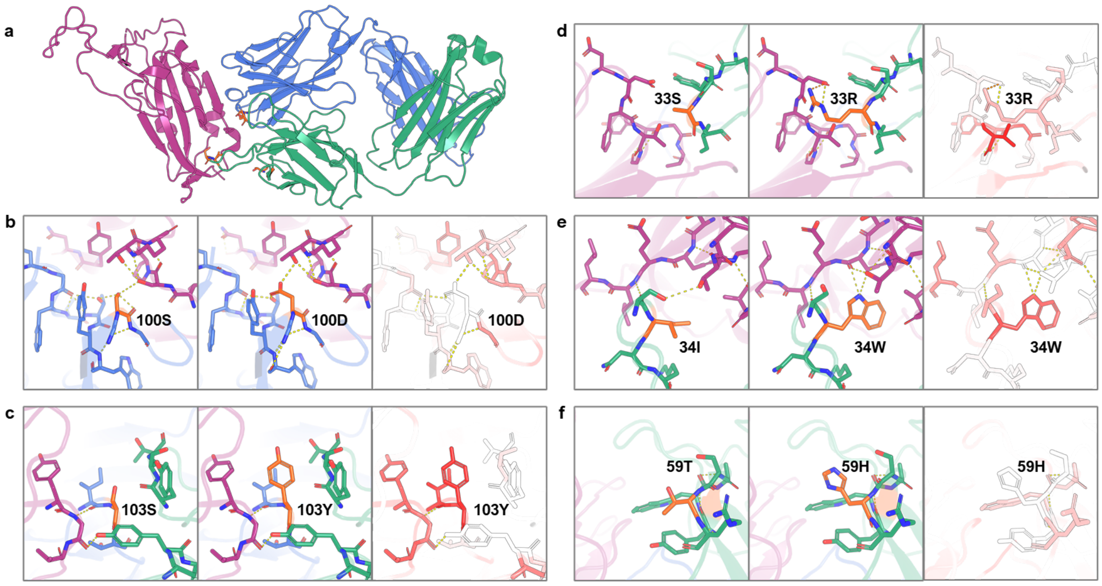


**Fig. S24.** Model attribution study of top mutations with enhanced binding effect in CR3022. (a) Overview of all 5 effective mutations sites predicted by GearBind. Figures (b-f) present triptych comparisons between wild type structures (left), mutant structures (middle), and the GearBind-predicted ΔΔ*G* contribution of each residue in mutant structure (right). Darker red denotes higher contributions. Specific mutations include: (b) S100D, (c) S103Y in the heavy chain, and (d) SL33R, (e) IL34W, (f) TL59H in the light chain. Structures presented here are modelled using Flex-ddG.

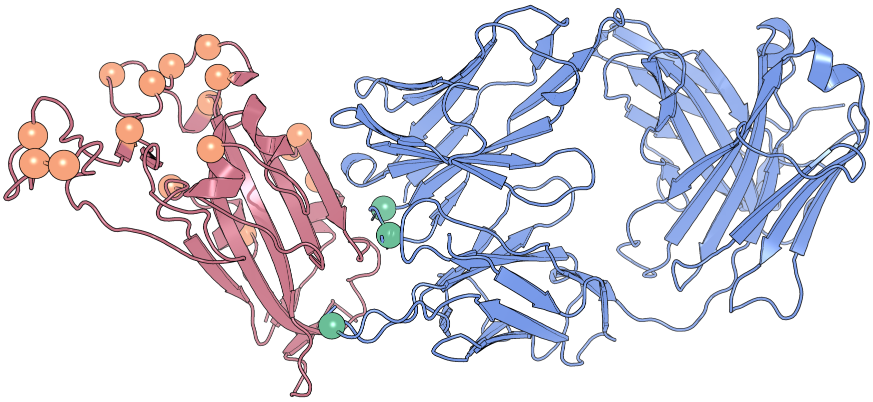

**Fig. S25.** Visualization of the locations of Omicron mutations (orange) and the mutation sites of the GearBind-designed triple mutant SH100D+SH103Y+SL33R (green). The SH100D+SH103Y+SL33R mutation sites are distant from most Omicron mutations.

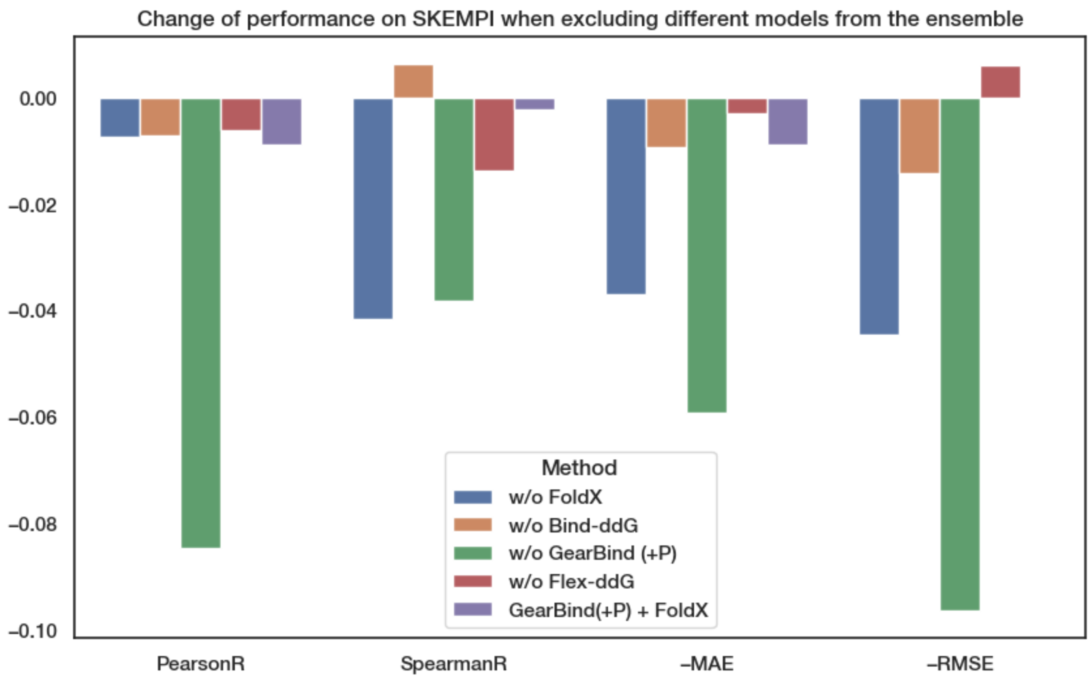


**Fig. S26.** Change of performance on SKEMPI (*n* = 5729) when excluding different models from the FoldX + Flex-ddG + Bind-ddG + GearBind (+P) ensemble. Note that the RMSE change of GearBind(+P) + FoldX compared to the whole ensemble is 0.000, making it invisible on the plot.

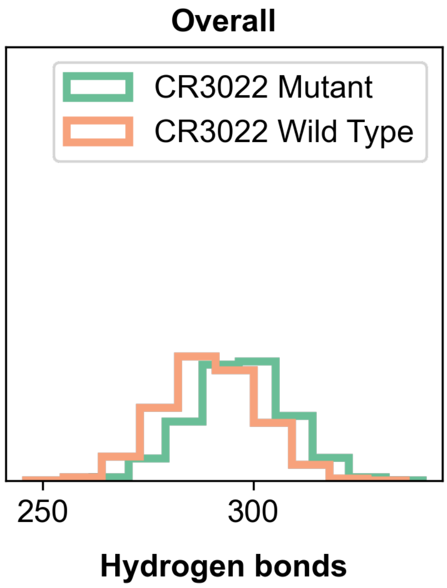


**Fig. S27.** The number of overall hydrogen bonds in the molecular dynamics simulation of the SARS-CoV-2 RBD-CR3022 complex.

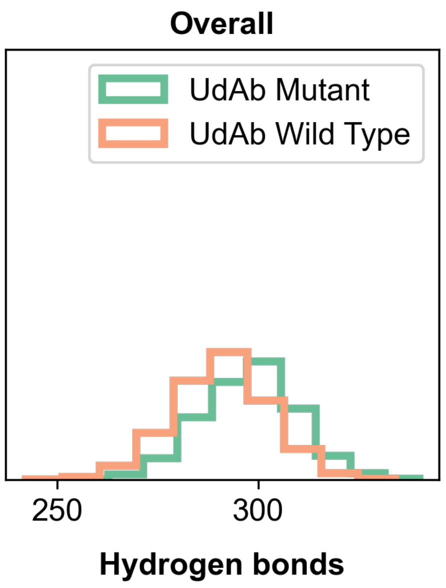


**Fig. S28.** The number of overall hydrogen bonds in the molecular dynamics simulation of the 5T4-UdAb complex.

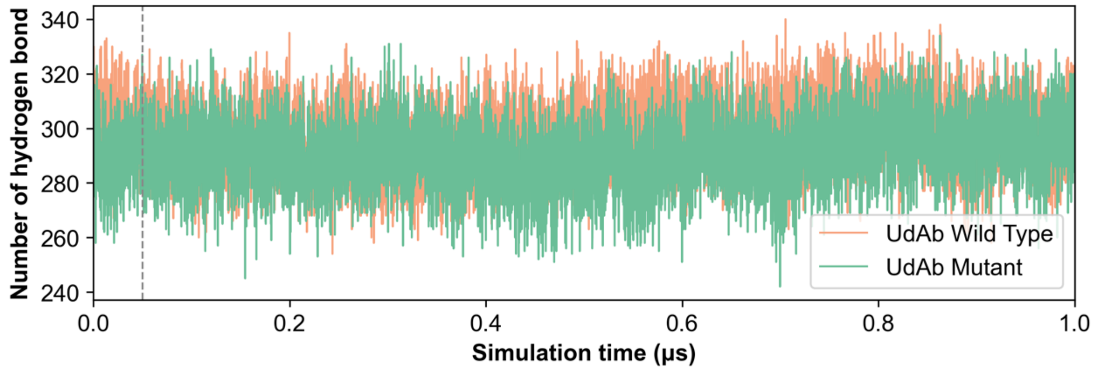


**Fig. S29.** The number of overall hydrogen bonds in the molecular dynamics simulation of CR3022 and RBD complex over simulation time**.**

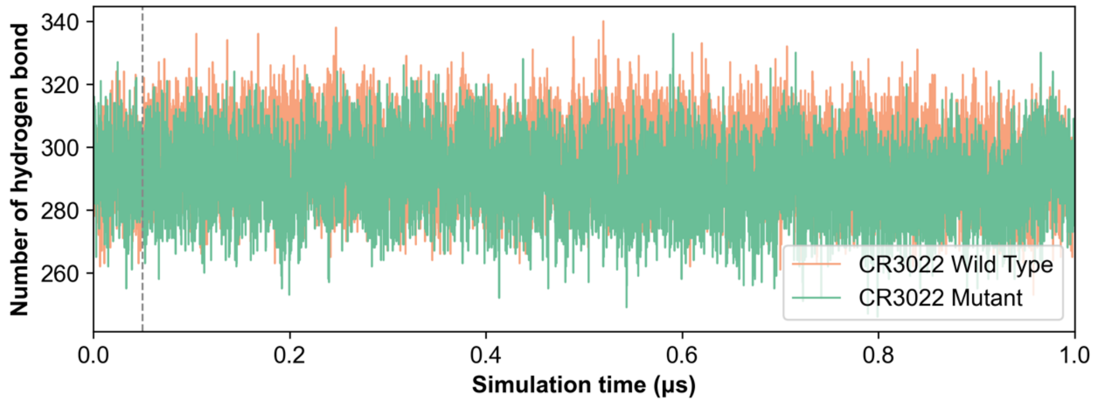


**Fig. S30.** The number of overall hydrogen bonds in the molecular dynamics simulation of UdAb and 5T4 complex over simulation time.
